# Sequencing and chromosome-scale assembly of the giant *Pleurodeles waltl* genome

**DOI:** 10.1101/2022.10.19.512763

**Authors:** Thomas Brown, Ahmed Elewa, Svetlana Iarovenko, Elaiyaraja Subramanian, Alberto Joven Araus, Andreas Petzold, Miyuki Suzuki, Ken-ichi T. Suzuki, Toshinori Hayashi, Atsushi Toyoda, Catarina Oliveira, Ekaterina Osipova, Nicholas D. Leigh, Andras Simon, Maximina H. Yun

## Abstract

The Iberian ribbed newt (*Pleurodeles waltl*) constitutes a central model for probing the basis of vertebrate regeneration. Here, we present the sequencing and chromosome-scale assembly of the 20.3Gb *P. waltl* genome, which exhibits the highest level of contiguity and completeness among giant genome assemblies. We uncover that DNA transposable elements are the major contributors to its expansion, with hAT transposons comprising a large portion of repeats. Several hATs are actively transcribed and differentially expressed during adult *P. waltl* limb regeneration, along with domesticated hAT transposons of the ZBED transcription factor family. Despite its size, syntenic relationships are conserved. As an example, we show the high degree of conservation of the regeneration-associated Tig1 locus with several neighbouring genes. Together, the *P. waltl* genome provides a fundamental resource for the study of regenerative, developmental and evolutionary principles.

## Introduction

Uncovering the basis underlying the ability to regenerate complex structures, as manifested in a few exceptional animal species, remains a central goal in regenerative biology. Featuring the widest repertoire of regenerative abilities among vertebrates^1–4^, salamanders constitute ideal models to accomplish this aim. While an impressive range of research tools has been developed for salamander species, in particular axolotls and Iberian ribbed newts^4–9^, their exploitation has been hampered by a scarcity of genomic resources. This is largely attributed to their enormous genome size (ranging from 14 to 120Gb,^10^) and substantial enrichment in repetitive sequences, which represent significant hurdles for sequencing, assembly generation and annotation. Recent developments in long-read sequencing technologies^11^, assembly algorithms^12, 13^ and chromosome conformation capture methods (Hi-C)^14^ have enabled the generation of high-quality assemblies for species with giant genomes such as lungfish (c.40Gb, ^15, 16^) and axolotl (c. 32Gb, ^17–19^). Comparisons between axolotl and newts have identified key differences in terms of life cycle, regeneration repertoire, regeneration mechanisms and genome composition ^2, 20^. Hence, it is important to characterize newt genomes in detail, to reveal the genetic basis of salamander-as well as species-specific mechanisms and evolutionary innovations. Among newts, *Pleurodeles waltl (P. waltl)* has emerged as an ideal laboratory model because of its experimental tractability^6, 7^. Prior attempts to sequence its giant genome resulted in an assembly based on Illumina short-reads^5^, providing the first source of genomic information for *P. waltl* albeit with limited completeness and contiguity. Here, we leveraged advanced long-read and chromosome conformation capture technologies to generate a highly-contiguous, complete chromosome-scale assembly for the giant genome of the Iberian ribbed newt. Further, we illustrate the applicability of this resource by revealing several attributes of the *P. waltl* genome of relevance to regeneration research and comparative genomics.

## Results and discussion

### Sequencing, chromosome-scale assembly and annotation of the *Pleurodeles waltl* genome

In order to overcome the challenges inherent to sequencing and assembling a giant genome, we took advantage of the highly-accurate Pacific Biosciences (PacBio) HiFi sequencing (Fig. 1a). We sequenced genomic DNA from a female *P. waltl* newt (Supplementary Fig. 1) generating 44,683,779 long-reads, representing 41X genome coverage. We employed *HiFiasm*^12^ and purge-dups^21^ yielding a 20.3Gb contig assembly of remarkably high contiguity with an N50 of 45.6Mb and N90 of 11.1Mb (Fig. 1b). Indeed, its contiguity is increased by an order of magnitude compared to the *Protopterus annectens* assembly, based on Oxford nanopore^16^, two orders of magnitude compared to the *Ambystoma mexicanum* assembly, based on PacBio RSII reads^18^, and 4 orders of magnitude compared to the short-read *P. waltl* assembly ^5^ (Fig. 1c). Kmer-based analysis indicated a level of heterozygosity of 0.39% (Supplementary data 1).

**Fig. 1.**
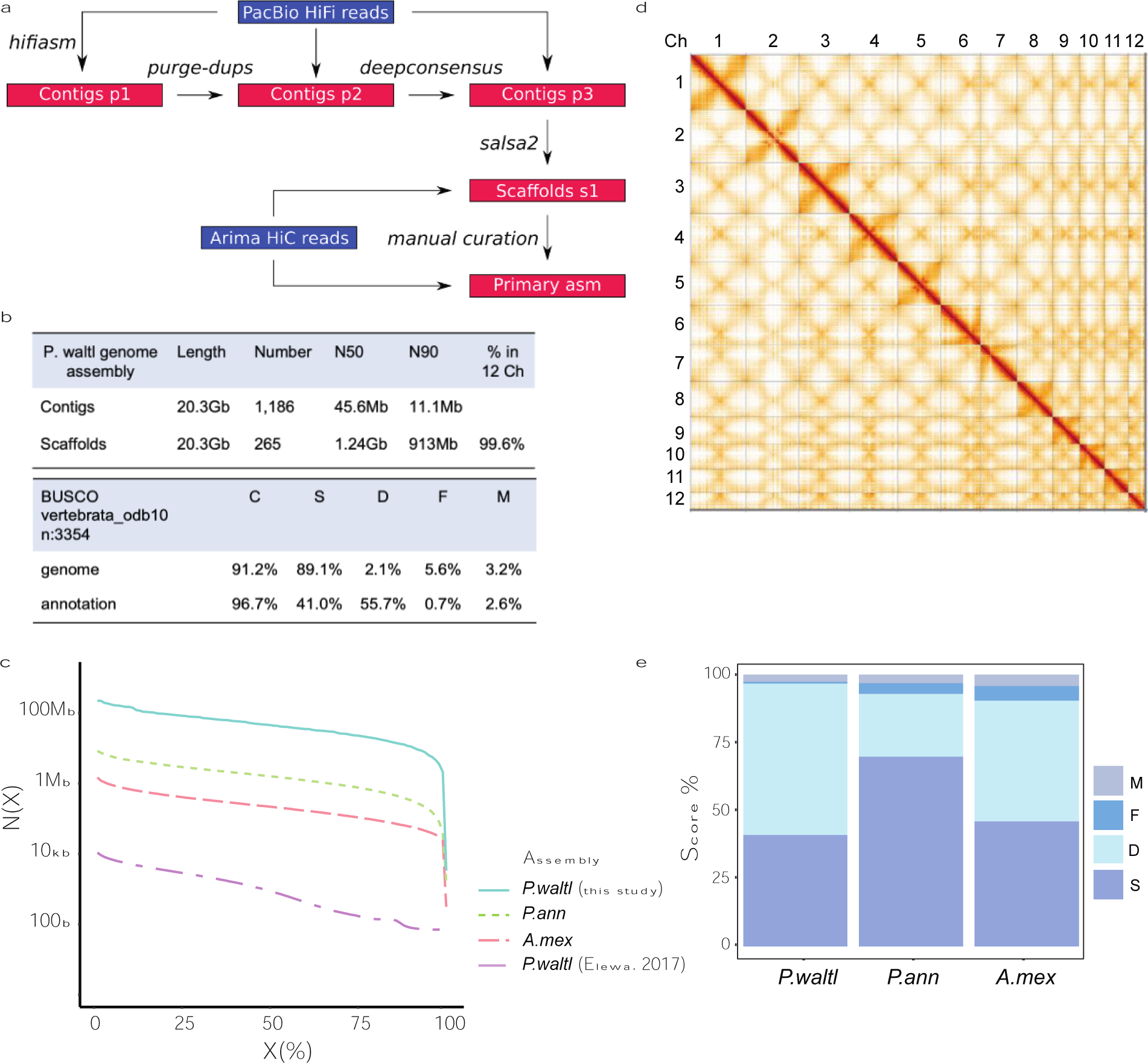
Sequencing and chromosome-level assembly of the *P. waltl* genome. (a) Schematic representation of the sequencing and assembly strategy. (b) *P. waltl* genome assembly features (top) and BUSCO assessment (bottom). (c) Contig N(X) plot showing which % of each assembled genome (X) is contained within pieces at least N(X) bp in size. Shown are contig statistics from *Pleurodeles waltl* (this study and ^5^), *Protopterus annectens*^16^ and *Ambystoma mexicanum*^18^. (d) Hi-C interaction heatmap of contact data for scaffolded genome. Individual scaffolds are delineated. Denser areas of red signal off-diagonal represent interactions between the arms of the same chromosome. (e) Gene completeness based on BUSCO single-copy vertebrata orthologs (n=3,354). Scores are based on annotations for *Pleurodeles waltl*, *Protopterus annectens* and *Ambystoma mexicanum*. C: Complete, S: Single copy, D: Duplicated, F: Fragmented, M: Missing.

Next, we set out to generate a high-resolution chromosome-scale assembly based on chromosome conformation capture (Hi-C) Illumina reads. We generated two Arima Hi-C libraries from liver tissue (6 billion read pairs - 1.8Tbp) of the same individual sequenced above (Supplementary Fig. 1). Following scaffolding with *salsa2* ^22^ and manual curation (Fig. 1a, Supplementary Fig. 2), we obtained a chromosome-level assembly with a scaffold N50 of 1.24Gb (chromosomes 1-4 are in two parts due to technical limitations of standard tools in processing scaffolds larger than 2Gb) in which 99.6% of the contigs could be assigned to the 12 chromosomes (Fig. 1b). The percentage of chromosome incorporation of contigs is on a par with the highest among all reported giant genomes at 99.6%^16^. The number and length of the chromosomes is in agreement with the reported *P. waltl* karyotype^5^. At the structural level, Hi-C contact analysis (Fig. 1d) and subsequent haplotype-phased assemblies (Supplementary Fig. 3) indicates the existence of two inversions between the two haplotypes of the sequenced individual, one in the central region of chromosome 2 (Supplementary Fig. 4, Supplementary data 2) and another one in chromosome 5 (Supplementary Fig. 5, Supplementary data 3).

To facilitate the annotation of genomic loci we took advantage of the PacBio Iso-seq platform, and generated full-length mRNA sequences for brain (1Gb,^23^) and spleen (0.86Gb) of the same newt used for sequencing. These were combined with PacBio Iso-seq data from adult limb blastema (1.76Gb), *de-novo* transcriptomes^5, 24^, and Augustus predictions. We identified a total of 18,799 conserved protein coding genes in our assembly, comprising 96.7% of vertebrate single-copy orthologs based on BUSCO analysis, with only 2.6% missing (Fig 1b). Gene completeness analysis based on BUSCO single-copy orthologs within vertebrata confirmed the high completeness of the *P. waltl* assembly in comparison to other giant genomes (Fig. 1e). Further, we found that *P. waltl* has a similar number of protein-coding genes compared to other vertebrates (Fig. 1e), indicating that whole-genome duplication events are not responsible for the expansion of the *P. waltl* genome.

### Transposable elements underlie *Pleurodeles waltl* genome expansion

Proliferation of transposable elements (TE) has emerged as a chief mechanism underlying the expansion of giant genomes, as observed in lungfish and axolotl^15–17^. Indeed, expansion of Gypsy retrotransposons and DNA Harbinger TEs were proposed to account for two thirds of the repetitive content based on the short-read *P. waltl* assembly^5^. We therefore performed repeat masking of our highly complete genome assembly, uncovering that 74% of the genome (corresponding to 15Gb) is made up of repetitive elements (Fig. 2a). Repeat content is higher than that of the giant axolotl genome based on our own (68%, Fig. 2a) and previously reported analyses^18^. This may be attributable to either a higher retention of transposable elements within the Iberian newt genome, or the higher contiguity and completeness of its assembly. Further, the percentage of repetitive sequences in the Iberian ribbed newt genome is comparable to that of lungfish^15, 16^, which exhibits the highest repetitive content found in the animal kingdom. Unlike in the axolotl, where long terminal repeat (LTR) transposable elements are dominant (45% of repeats, 8.7Gb), the DNA repeat class is the major contributor (51% of repeats, 7.6Gb) to the *P. waltl* genome (Fig. 2a). In depth analysis of the genomic contributions of each repeat superfamily (Fig. 2b) revealed that Gypsy (19.7% of repeats, 3Gb) and Harbinger (19.6% of repeats, 2.9Gb) elements are among the most represented superfamilies, as previously suggested^5^. Notably, it also uncovered a large contribution of DNA/hAT (hobo-Ac-Tam3) transposable elements (15% of repeats, 2.3Gb) to the *P. waltl* genome. Among the predominant DNA/hAT elements, the non-autonomous Miniature Inverted Repeat Transposable Element (MITE) and nMITE types make up 15% of all repeats in the assembly (Fig. 2c).

**Fig. 2.**
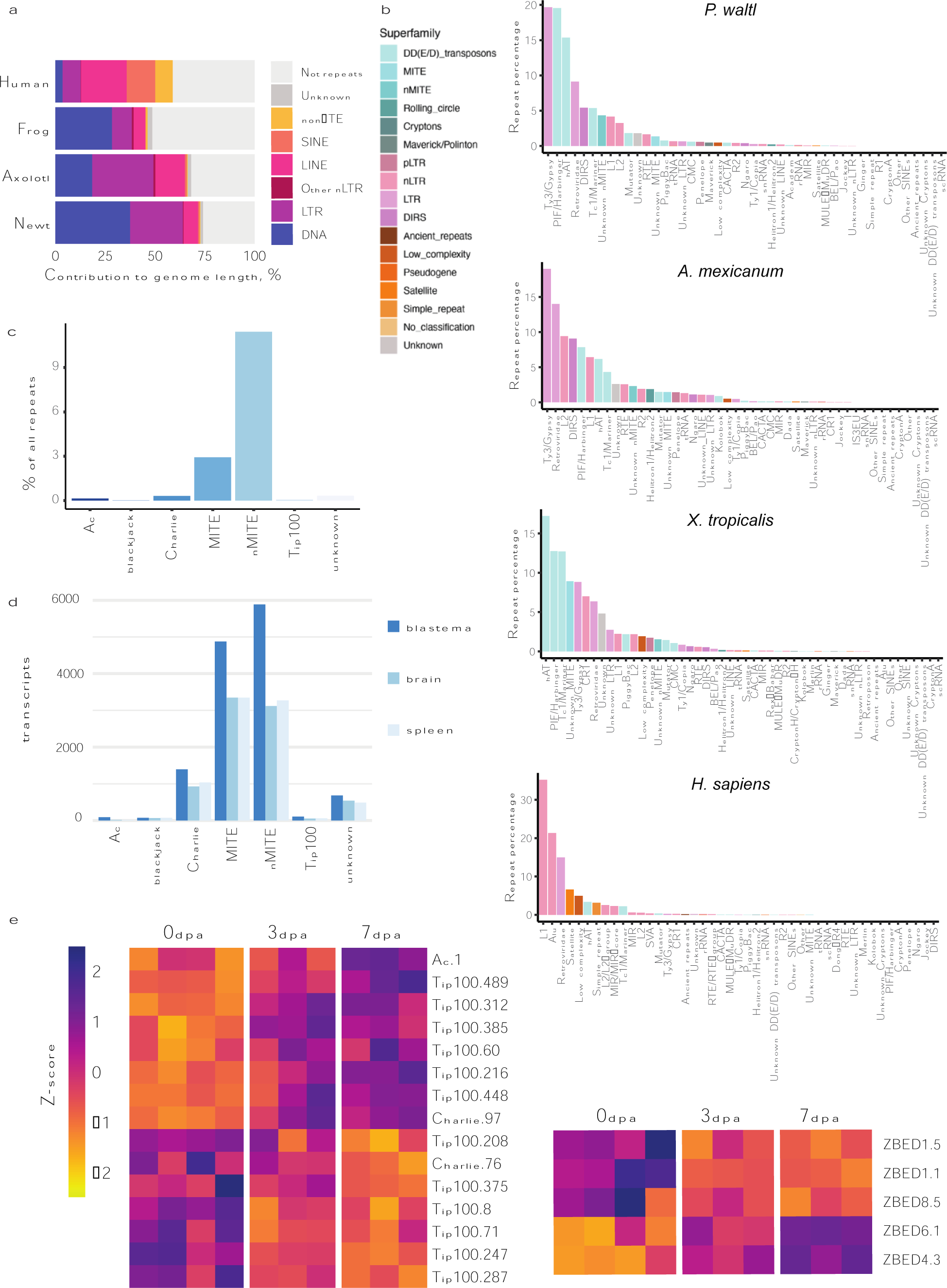
Genome expansion, repeat element composition and regeneration-associated expression of hATs and domesticated hATs in *P. waltl*. (a) Pie charts showing the contribution of the main repeat element (RE) classes to each of the indicated genomes. Note the abundance of DNA repeat elements in *P. waltl*. (b) Contribution of repeat element superfamilies to each indicated genome. (c) Relative contribution of hAT families to the total RE repertoire of the *P. waltl* genome, expressed as percentage of all RE based on sequence length. (d) Expression of indicated hAT elements in PacBio Iso-seq-derived transcriptomes from *P. waltl* limb blastema, brain and spleen. (e) Differential expression of hATs (left) as well as domesticated hATs (ZBED1, ZBED4, ZBED6, ZBED8) and corresponding isoforms (right) during limb regeneration in *P. waltl*, based on normalised and centered RNAseq counts for the indicated conditions. Only genes whose differential expression is significantly altered (between 0 and 3dpa, or 0 and 7dpa) are depicted. Colour key represents Z-score values.

With regards to both repeat element class and superfamily contributions (Fig. 2a,b), significant differences arise among representative vertebrate genomes, even between the Iberian newt and the axolotl, indicating that independent mechanisms drive genome expansion in each organism. To gain insights into the expansion history of transposable elements in the *P. waltl* genome, we performed Kimura distance analysis, whereby the number of substitutions to the consensus sequence of each element is used to estimate its relative age. Results highlight differential expansion kinetics between the main transposable element types (Supplementary Fig. 6a,b), with DNA/hAT and Harbinger elements having undergone one wave of expansion with overlapping periods, and LTR/Gypsy elements exhibiting a longer period of basal activity, including a very recent, extended wave of expansion (Supplementary Fig. 6). Thus, the *P. waltl* genome has sustained several waves of TE-driven expansion.

Bursts of transposition are key sculptors of the genome landscape. For example, DNA/hAT elements are distributed primarily within intergenic regions (76%, Supplementary Fig. 7) yet they are also found in exons (0.07%), contributing to the exapted or ‘domesticated hAT’ gene category, and are abundant among introns (23.85%). This distribution resembles that observed for all repeat elements within the *P. waltl* genome (Supplementary Fig. 7). Transposable element contributions have resulted in an increase in intron dimensions in *P. waltl* (Supplementary Fig. 8), leading to a median (6Kb), mean (17.8Kb) and maximum (4.8Mb) intron size several orders of magnitude greater than in frogs, mice and humans^17^. Intergenic regions in the *P. waltl* genome also displayed a significant expansion, with a median of 50Kb and a mean of 116Kb. In contrast, *P. waltl* exon dimensions (median: 159bp; mean: 277bp; maximum: 200Kb) are comparable to other vertebrates (Supplementary Fig. 7). These data show that transpositions shape gene structure and may impact gene function in *P. waltl*.

### DNA/hAT and domesticated hAT expression during *Pleurodeles waltl* limb regeneration

The expansion of DNA/hAT elements prompted us to probe whether these were being expressed. Mining of the PacBio Iso-seq transcriptomes from *P. waltl* adult limb blastema, brain and spleen (Fig. 2d) indicate that all hAT types are expressed in these tissues, albeit to a different degree. Notably, these include several hATs longer than 2000bp, thus likely to contain full transposition elements (Supplementary Fig. 9).

We further observed that the diversity of expressed hAT elements (Supplementary Fig. 9) and number of transcribed copies (Supplementary Fig. 10) is higher in limb blastema than in the other tissues examined. Thus, we went on to probe available RNAseq datasets of *P. waltl* limb regeneration stages^5^. Differential expression analysis uncovered dynamic transcriptional changes in several Tip100, Charlie, MITE and Ac hATs during limb regeneration (Fig. 2e, Supplementary data 4), both after wounding (3 days post-amputation (dpa)) and at early blastema formation stages (7dpa). Further, we found that isoforms of various Zinc Finger BED-type (ZBED) gene family members, which derive from hAT transposons following a ‘domestication’ event^25^, are upregulated during limb regeneration (Fig. 2e, Supplementary data 5). These include ZBED6, a transcription factor involved in muscle development^26^, ZBED1, a putative transcriptional regulator of cell proliferation and ribosomal gene expression^27^, and ZBED4, a retina-associated factor^28^, isoforms of which are found significantly upregulated during blastema formation. The latter is of particular interest in light of a previous report of ZBED4 upregulation in lungfish tail blastemas^29^. A number of ZBED isoforms are also differentially expressed across additional *P. waltl* tissues, including in the germline (Supplementary Fig. 11). The existence of several isoforms for each ZBED gene is worthy of note, as they may be indicative of recent transposition events.

To further investigate the expression of hATs and domesticated hATs in an *in vivo* setting, we analysed the expression of two ZBED family members (ZBED1 and ZBED6) and a representative hAT (Charlie 76) in mature limb and upper-arm blastema tissues at various limb regeneration stages (Fig. 3). Of note, the changes in gene expression between mature limb and early blastema for all three elements recapitulated the pattern observed in the transcriptome analysis (Fig. 2e). Similar observations were obtained using an alternative normalizer (Supplementary Fig. 12). Collectively, these data show that, unlike in other vertebrate genomes^30, 31^, hATs are actively transcribed and thus may continue to shape the *P. waltl* genomic landscape. Further, they suggest a potential functional involvement of hATs and domesticated hATs in limb regeneration, an avenue for further exploration.

**Fig. 3.**
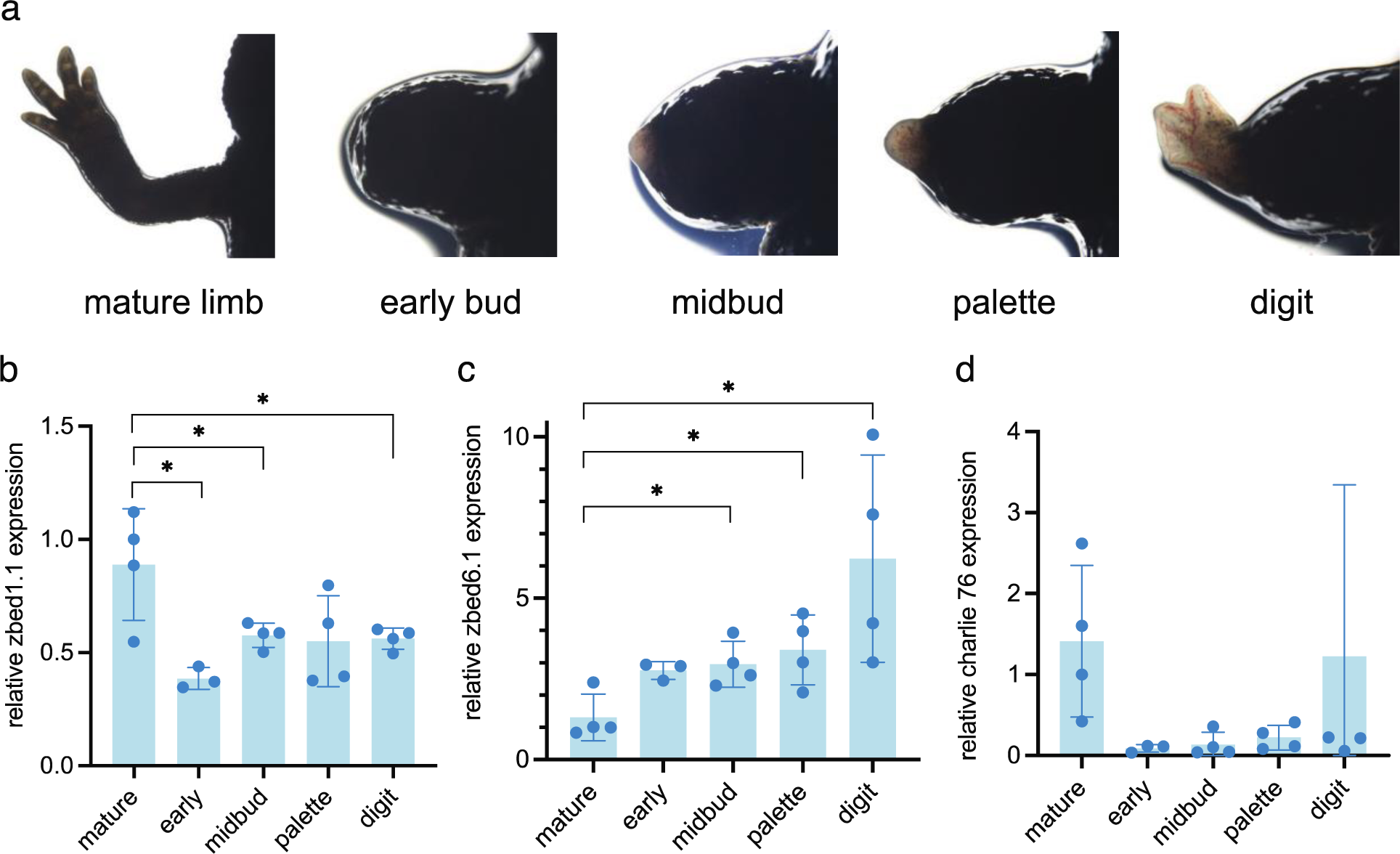
Relative expression of classical and domesticated hATs isoforms during *P. waltl* limb regeneration. **(a)** Representative images of selected stages during limb regeneration. (b) qRT-PCR quantification of relative gene expression for the indicated domesticated (zbed1 (b), zbed6 (c)) and classical hAT (charlie 76 (d)) isoforms. *actb2* was used as a normaliser. *p < 0.05 (Welch ANOVA test followed by t-test individual comparison). Error bars represent SD.

### Synteny conservation

The gigantic TE-dependent expansion observed in the *P. waltl* genome could have led to the disruption of syntenic relationships between *P. waltl* and vertebrate orthologues. To address this, we first performed macrosynteny comparisons of 1:1 orthologous genes between *P. waltl* and representative jawed vertebrates (phylogenetic relationship shown in Fig. 4a). Our analysis revealed a high level of syntenic conservation based on 6,434 1:1 orthologues between the Iberian ribbed newt and the axolotl assemblies (Fig. 4b) as well as the local organisation of 11,925 orthologues found in the Gar genome (Fig. 4c), identifying homologous chromosomal tracts between species. Conservation of macrosyntenic correspondence is observed for additional regenerative organisms such as *Xenopus* (Supplementary Fig. 13) and lungfish (Supplementary Fig. 14), as well as for human (Supplementary Fig. 15). Of note, the analysis of syntenic boundaries between *P. waltl* and axolotl as well as *Xenopus* provide independent evidence for the aforementioned inversion of chromosome 5 in one haplotype, indicated by the Hi-C data (Supplementary Fig. 5, 13). Altogether, this analysis provides further proof of the quality of the chromosome-scale assembly, and offers insights into its macrosyntenic conservation.

**Fig. 4.**
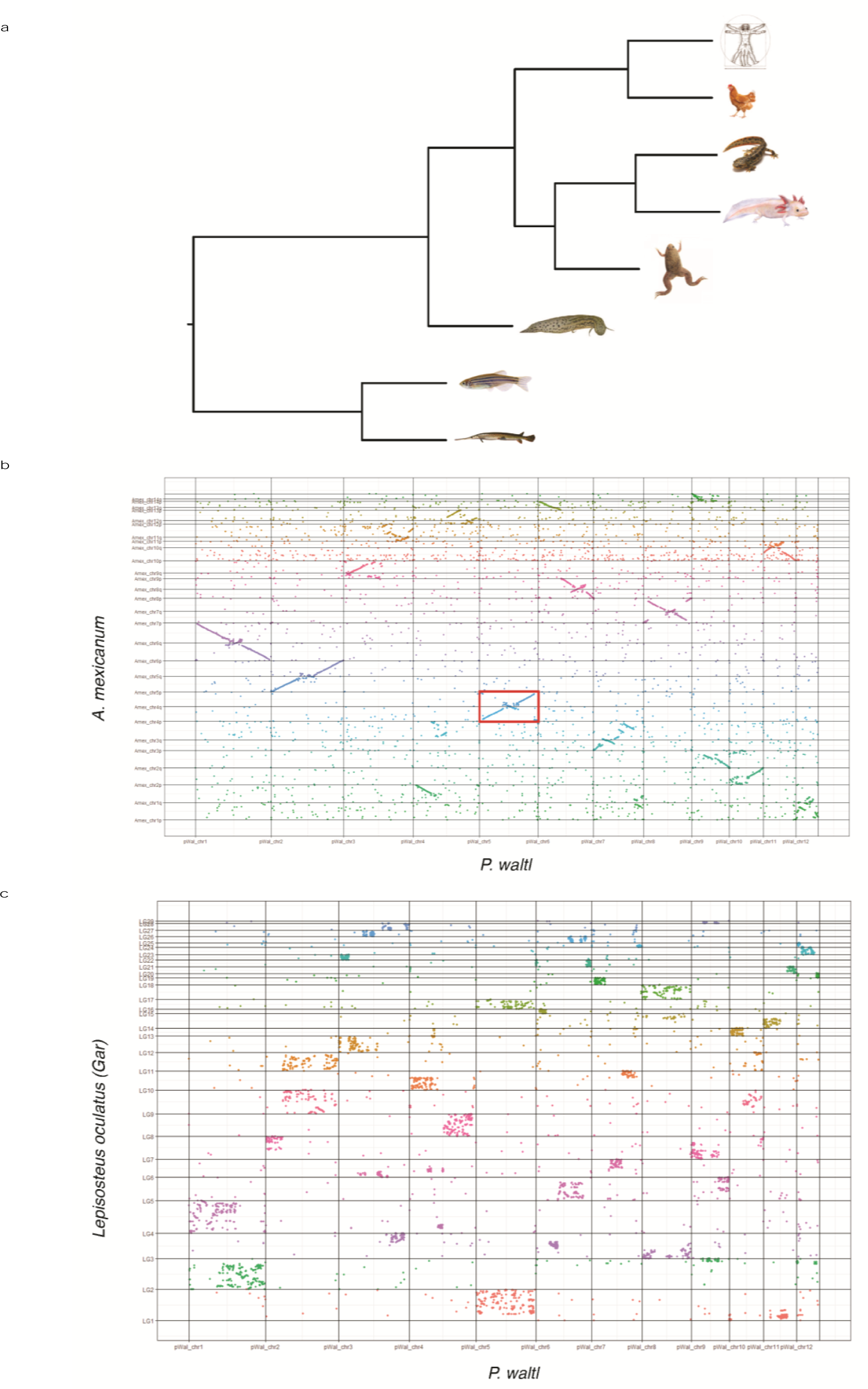
Comparative macrosyteny analysis of the *P. waltl* genome. (a) Phylogenetic relationships between the indicated species constructed using the maximum-pseudolikelihood coalescent method MP-EST, based on one-to-one orthologues identified through blastp between *Pleurodeles waltl* and *Ambystoma mexicanum*, *Xenopus tropicalis*, *Lepisosteus oculatus*, *Danio rerio*, *Protopterus annectens*, *Homo sapiens* and *Gallus gallus*. Vitruvian man: © A. Prahlad. (b) Oxford plot of macrosyntenic relationships between *P. waltl* and *A. mexicanum* chromosomes. Coloured dots indicate relative chromosomal arrangement of newt-axolotl orthologues. Inversion of *P. waltl* chromosome 5 is indicated by the red rectangle. (c) Oxford plot depicting macrosyntenic relationships between *P. waltl* and Gar (*L. oculatus*) orthologues. LG: linkage group, chr: chromosome.

To probe synteny conservation at the gene loci level, we next centered on the *Tig1*/*Rarres1* locus (Fig. 5a, see also Supplementary data S6 for a general pipeline for microsynteny analysis), recently implicated in the regulation of proximo-distal identity during salamander limb regeneration^32^. Despite clear variations in gene size, we uncovered a remarkable conservation of loci organisation, with the presence of *Tig1* neighbours *Schip1* and *Mfsd1* at the 5’ end, and of *Gfm1*, *Mlf1* and *Shox2* at the 3’ end of the *Tig1* locus, a structure that is recapitulated across representative vertebrate genomes (Fig. 5a). Within this exemplary region of the *P. waltl* genome, the major contributor to gene size change is intron expansion, driven by accumulation of TEs such as hATs, Gypsy and Harbinger (Fig. 5b). Intron size differs for each orthologous gene across species, indicative of independent expansion events (Fig. 5a). In agreement, we observed differential contribution of transposon elements to intron size in genes within the locus (Fig. 5b). While most of the aforementioned genes seem to have undergone significant expansion during evolution, the orthologous of *Shox2,* a gene involved in limb development^33^ remains stable, displaying a lack of intron expansion in all species analysed. This is consistent with the notion that intron size of key developmental genes is under constraint in evolution, possibly related to their transcriptional requirements^17^.

**Fig. 5.**
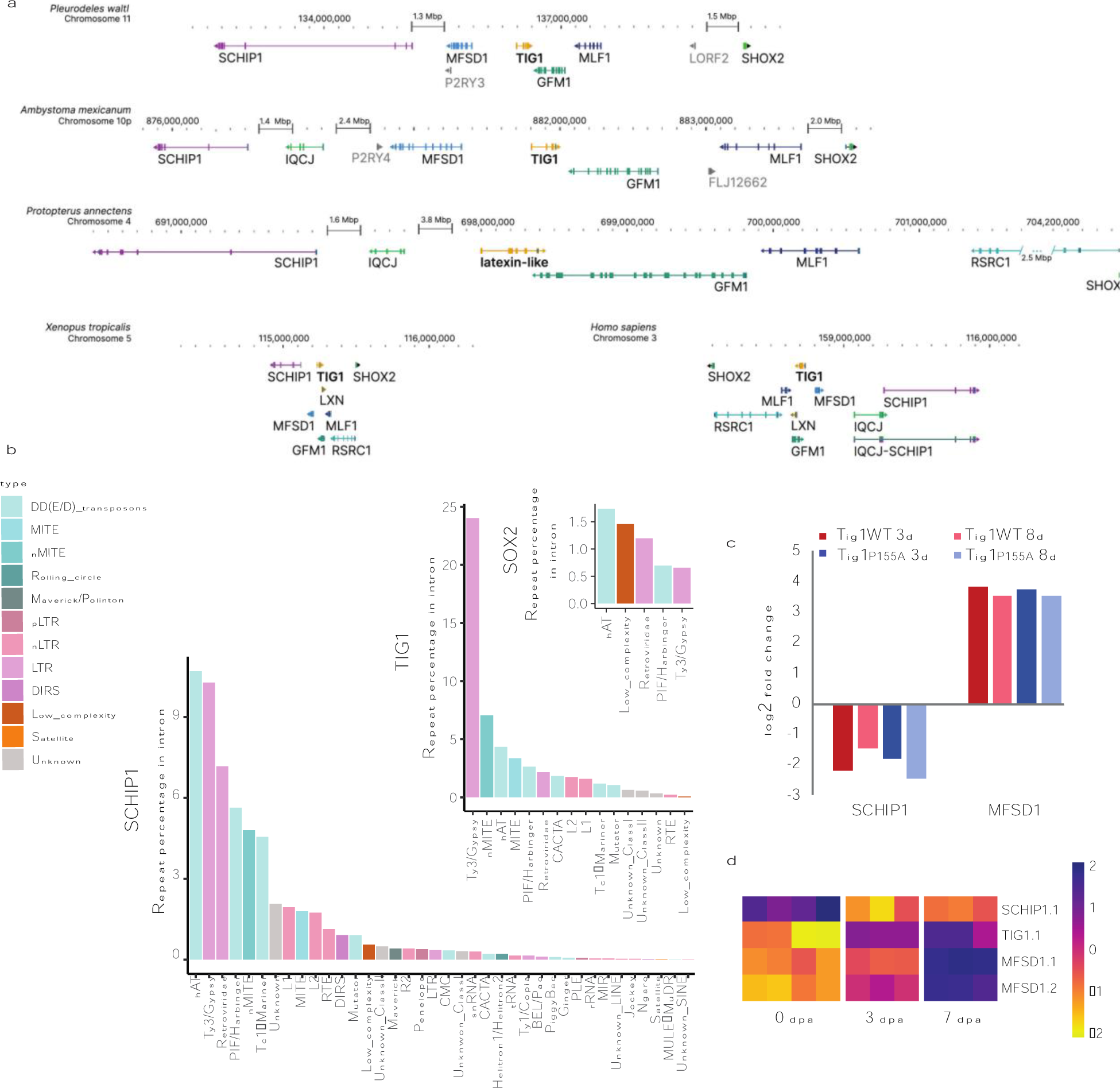
Microsynteny conservation of the *Tig1* locus. (a) Analysis of genomic location and structure of genes associated with the *Tig1* locus in the indicated tetrapod species. Genes are indicated in colour. Arrowheads indicate reading frame direction. Perpendicular bars within genes represent exons. Note the increase in intron size for all genes associated with the *Tig1* locus in *P. waltl* compared to *X. laevis* and *H. sapiens*. Variation in intron length among species with giant genomes is indicative of independent intron expansion. (b) Relative contribution of repeat elements to intron expansion for the indicated genes. (c) Tig1 overexpression affects expression of nearby genes in *A. mexicanum*. Bars represent ratios of gene expression in Tig1 or Tig1^P155A^ overexpressing cells versus control cells (analysis based on dataset from ^32^). (d) Heatmap depicting differences in gene expression of Tig1 and its genomic neighbours during *P. waltl* limb regeneration.

The conservation of the *Tig1*-*Mfsd1-Schip1* arrangement might bear functional implications. Both *Schip1* and *Mfsd1* are differentially regulated during Tig1-mediated reprogramming of distal cells to a proximal identity in axolotls (Fig. 5c), with *Mfsd1* upregulated and *Schip1* downregulated upon Tig1 overexpression in distal cells. Similar regulatory effects are observed upon reprogramming with a hyper-proximalising version, Tig1^P155A^ (Fig. 5c). Accordingly, the three genes are differentially regulated during limb regeneration in *P. waltl* (Fig. 5d, Supplementary data 7) in a manner that reflects the proposed hierarchy. Therefore, the preservation of their genomic location relative to *Tig1* could result from their participation in a common regulatory network.

### Concluding remarks

The genomic and transcriptomic resources hereby presented provide new insights into the evolution of giant genomes and the molecular underpinning of complex regeneration programmes. The chromosome scale assembly for the Iberian ribbed newt exhibits the highest contiguity and completeness among all giant genomes reported to date^15, 16, 18^. DNA transposable elements are the major contributors to the genome size, and uncover a significant role played by hAT elements in genome expansion and the shaping of gene loci architecture. Moreover, their functional impact may extend beyond the latter, as we find that many hATs are actively transcribed in various tissues, and members of this transposon superfamily as well as hAT-derived genes are regulated during limb regeneration in adult newts. Hence, hATs and related members of the family may represent a previously overlooked set of contributors to salamander limb regeneration.

Despite its size, the *P. waltl* genome exhibits a striking conservation of syntenic relationships at both macro- and micro-levels. In the case of the proximo-distal determinant Tig1, our study suggests that its genomic arrangement reflects a regulatory connection between Tig1 and the genes it clusters with, of potential relevance to the remarkable regenerative abilities found in *P. waltl*.

Collectively, our findings highlight the usefulness of the *P. waltl* chromosome-scale assembly as a resource for enhancing gene discovery in the context of development, physiology and regeneration, while providing a foundation for epigenetic studies, enhanced gene-editing and the detailed exploration of regulatory networks underlying biological processes.

Genome and annotation files are available through the Max Planck Digital Library (https://doi.org/10.17617/3.90C1ND) and through NCBI under BioProject PRJNA847026.

## Materials and Methods

### Animal procedures

All animal experiments were performed in accordance with the guidelines of local, European and Japanese ethical permits. Adult Iberian Ribbed newts (*Pleurodeles waltl*) were obtained from the aquatic animal facility in Karolinska Institutet (genome sequencing) and CRTD-Center for Regenerative Therapies Dresden (qRT-PCR analysis), where they were bred, raised and kept in captivity for generations as described^34^. For genome sequencing, only female newts were used, as in this species the females represent the sex that display a heterogametic ZW genotype involved in sex determination^35^. To ensure that sequencing would be performed in a diploid animal, we analysed tail tip tissue from a tail clip from six adult females. The tissue collected was fixed, processed, sectioned at 30µm, and stained with DAPI to highlight the DNA according to^36^. The 2n DNA content was confirmed by analysing both the expected size of the nuclei as well as chromosome quantification in cartilage cells found in the M-phase of the cell cycle (2n = 24 chromosomes, ^5^) (Supplementary Fig. 1). A 14.1cm-long adult female weighing 17.93g was selected for tissue collection. Tissues from the same animal were used for genome sequencing, Hi-C as well as brain and spleen Iso-Seq. After the tail clip was allowed to regenerate, the animal was deeply anaesthetised in 0.1% tricaine (MS-222, Sigma, pH=7) and sacrificed by decapitation. Immediately after, tissue dissection and collection were performed by three researchers working simultaneously to minimise sample degradation. The samples collected consisted of blood, brain, spinal cord, limb skeletal muscle, spleen, heart, and liver. Samples were snap-frozen in liquid nitrogen and stored at −80°C for further processing.

For blastema Iso-seq preparation, adult newts were obtained from the Amphibian research center at Hiroshima University, Japan, and their rearing and treatments were performed and approved in accordance with the Guidelines for the Use and Care of Experimental Animals and the Institutional Animal Care and Use Committee of National Institute for Basic Biology, Japan.

For qRT-PCR analysis of the indicated limb regeneration stages, newts were anesthetised in 0.03% benzocaine (Sigma) prior to limb amputation at the mid-humerus level. Animals were allowed to recover and regenerate at 20 °C, and limbs or blastemas collected and processed for RNA extraction as specified below. Experiments were performed in accordance with the laws and regulations of the State of Saxony.

### Genome sequencing

The spleen from a female, adult *Pleurodeles waltl* was collected. Genomic DNA from the spleen was extracted using the Monarch® HMW DNA Extraction Kit for Tissue (NEB #T3060) as per manufacturer’s protocol. DNA was then immediately frozen and stored at −20°C until further downstream processing. Input QC of the DNA was performed using Dropsense, Qubit and Femto pulse to evaluate concentration, purity and size. Sample libraries were prepared according to Pacbio’s Procedure & Checklist – Preparing HiFi SMRTbell® Libraries using the SMRTbell Express Template Prep Kit 2.0, PN 101-853-100 Version 05 (August 2021) using the SMRTbell Express Template Prep Kit 2.0. Samples were sheared on Megaruptor 3 with speed setting 31 followed by 32. An Ampure bead purification was performed after the shearing. The samples were size selected using SageElf, according to Pacbio’s protocol. Fractions 1-4 were used for sequencing. Quality control of sheared DNA and SMRTbell libraries was performed on Fragment analyzer, using the Large Fragment standard sensitivity 492 kit. Primer annealing and polymerase binding was performed using the Sequel II binding kit 2.2. They were sequenced on the Sequel II and IIe instrument, using the Sequel II sequencing plate 2.0 and the Sequel® II SMRT® Cell 8M, movie time 30 hours and pre-extension time 2 hours.

### Hi-C

Chromatin conformation capturing was performed using the ARIMA-HiC+ High Coverage Kit (Article Nr. A101030-ARI) following the user guide for animal tissues (ARIMA Document, Part Number: A160162 v00). In brief, 40 mg flash-frozen powdered heart tissue was chemically crosslinked. The crosslinked genomic DNA was digested with a restriction enzyme cocktail consisting of four restriction enzymes. The 5’-overhangs were filled in and labelled with biotin. Spatially proximal digested DNA ends were ligated and purified. The proximally-ligated DNA was then sheared and enriched for the ligated biotin containing fragments. Due to the expected genome size, two barcoded Illumina sequencing libraries were prepared following the ARIMA user guide for Library preparation using the Kapa Hyper Prep kit (ARIMA Document Part Number A160139 v00). The barcoded Hi-C libraries were sequenced on a NovaSeq6000 (NovaSeq Control Software 1.7.5/RTA v3.4.4) with a 151nt(Read1)-10nt(Index1)-10nt(Index2)-151nt(Read2) setup using ’NovaSeqXp’ workflow in ’S4’ mode flowcell. The Bcl to FastQ conversion was performed using bcl2fastq_v2.20.0.422 from the CASAVA software suite. The quality scale used was Sanger / phred33 / Illumina 1.8+.

### Iso-seq

Spleen Iso-seq: Tissue was harvested from the spleen of a female adult *Pleurodeles waltl*. Tissue was immediately frozen on dry ice and stored at −80 °C. Approximately 10 mg of each indicated tissue were pulverised in liquid nitrogen and used for RNA extraction by means of Total RNA Purification Kit (Cat. 17200 from Norgen Biotek) as per manufacturer’s instructions. The pulverised spleen powders are mixed well with the β-mercaptoethanol containing buffer RL and the homogenates are further passing through the needle attached syringe for 5-10 times. The genomic DNA was removed via on column DNA removal using Norgen RNAse free DNAase kit (Cat. 25710). Input QC of the RNA was performed on the Agilent Bioanalyzer instrument, using the Eukaryote Total RNA Nano kit to evaluate RIN and concentration. RIN value obtained was above 9 for both samples. The sample libraries were prepared according to Pacbio’s Procedure & Checklist – Iso-Seq™ Express Template Preparation for Sequel® and Sequel II Systems, PN 101-763-800 Version 02 (October 2019) using the NEBNext® Single Cell/Low Input cDNA Synthesis & Amplification Module, the Iso-Seq Express Oligo Kit, ProNex beads and the SMRTbell Express Template Prep Kit 2.0. 300 ng RNA was used as input material. The samples were amplified 12 cycles. In the purification of amplified cDNA the standard workflow was applied (sample is composed primarily of transcripts centered around 2 kb). Quality control of the SMRTbell libraries was performed with the Qubit dsDNA HS kit and the Agilent Bioanalyzer High Sensitivity kit. Primer annealing and polymerase binding was performed using the Sequel II binding kit 2.0. Libraries were sequenced on the Sequel IIe instrument, using the Sequel II sequencing plate 2.0 and the Sequel® II SMRT® Cell 8M, movie time=24 hours and pre-extension time=2 hours (1 SMRT cell per sample). Raw sequencing data was fed into IsoSeq3 (https://github.com/PacificBiosciences/IsoSeq), as per standard instructions, to generate fasta outputs of high-quality and low-quality reads. The high quality fasta output was taken for further use as part of the reference transcriptome.

Brain Iso-seq: the dataset is published^23^ and publicly available at NCBI (SRX16252717). Of note, the brain tissues correspond to the same individual described above.

Blastema Iso-seq: Adult newts (1-year-old) were anaesthetised in 0.1% MS-222, and the forelimbs were amputated with a surgical knife at the middle of the forearm level. Blastema was collected at the late-bud blastema stage^37^ under anaesthesia, and 4∼6 blastemas were pooled for each library. Total RNAs were purified using RNeasy mini kit (Qiagen) or NucleoSpin RNA Plus XS (Takara Bio) kit. Sample libraries were prepared according to the following PacBio Iso-Seq protocols: Iso-SeqTM Template Preparation for Sequel Systems and Iso-SeqTM Express Template Preparation for Sequel and Sequel II Systems. One of four blastema libraries was treated with gRNA/Cas9 ribonucleocomplex which targets 20 transcripts highly expressed in blastema to deplete abundant transcripts *in vitro* after the first bead purification step, another was treated with tyr Cas9 RNP to act as negative control, and the other two samples were untreated (Suzuki et al., unpublished). Libraries were sequenced on the Sequel I or II in National Institute for Genetics, Japan.

### Genome assembly and scaffolding

CCS reads (rq > 0.99) were called from the subreads.bam file using PacBio ccs (v6.0.0). Two contig assemblies were then created using hifiasm (v 0.16.0-r369) with arguments −l2 using the ccs reads as input. Purge-dups (v1.2.3) was then run on the primary contigs to create the set of contigs used for scaffolding. A further run of purge-dups on the combined assembly made up of the purged output from the primary contigs and the alt contigs from hifiasm created the alternate assembly.

To polish the assembled contigs from the primary assembly, all CCS reads were mapped to the contig assemblies using pbmm2 with arguments: --preset CCS -N 1 and variants were called using DeepVariant (v 1.2.0). Sites were filtered for those with ‘genotype 1/1’ to specify that all or nearly all reads support an alternative sequence at this position and a ’PASS’ filter value to specify that the site passed DeepVariant’s internal filters. Base errors were then corrected using bcftools consensus (v 1.12). Based on the QV values produced by merqury (v 1.0) this resulted in an assembly with QV 72.9 and QV 54.8 using PacBio CCS reads and Arima V2 Hi-C reads as a kmer database, respectively. The discrepancy between these values can be attributed to the fact the CCS reads were used to assemble the genome itself, therefore overestimate the accuracy of the genome, whereas the Hi-C reads are not uniform and contain coverage gaps, resulting in an under-estimate. The “true” value is likely to lie between these two values, but without an independent sequencing dataset we are unable to estimate the true value.

For scaffolding, SALSA2 and the VGP’s Arima mapping pipeline were run (https://github.com/VGP/vgp-assembly/blob/master/pipeline/salsa/arima_mapping_pipeline.sh). Briefly, Hi-C reads were mapped to the contigs using BWA-MEM (v 0.7.17-r1198-dirty).

Alignments were then filtered using the Arima filter_five_end.pl script (https://github.com/VGP/vgp-assembly/blob/master/pipeline/salsa/filter_five_end.pl) and removed potential PCR duplicates with Picard’s MarkDuplicates (v 2.22.6).

The resulting read-sorted bed file was used as input for SALSA2 (v 2.2). A number of manual curation rounds were then performed to correct scaffolding errors and to scaffold those contigs which were not automatically scaffolded into the 12 chromosomes. To this end, cooler (v 0.8.11) and HiGlass (v 2.1.11) were used to visually inspect the Hi-C maps and SeqKit (v 0.13.2) was used to re-arrange contigs and scaffolds into chromosome-level scaffolds.

To generate two haplotype-phased assemblies, we used hifiasm (v 0.16.1) with both PacBio CCS and HiC reads as input with arguments:

-l2 –h1 hic_R1.fastq.gz –h2 hic_R2.fastq.gz

and subsequently ran purge-dups (v1.2.3) separately on the two haplotypes as above. Each haplotype was then scaffolded and manually curated separately as above, with the exception that yahs (v1.1a) was used to scaffold instead of SALSA2.

#### Repeat masking

In order to mask the P.waltl genome a de-novo repeat library was first created using RepeatModeler (v 2.0.2) with argument -LTRStruct. The resulting pleurodeles-specific repeat library was combined with the Dfam repeat library for all ancestors of pleurodeles with famdb.py (v 0.4.2):

famdb.py -i Dfam.h5 families --ancestors -f fasta_name "pleurodeles waltl" > pleurodelesWaltl_repeatLibrary.fasta

The combined repeat library was then used to mask the genome using RepeatMasker (v 4.1.2):

RepeatMasker -gff -xsmall -e crossmatch -libdir RepeatMasker-4.1.2-p1/Libraries -lib aPleWal_combinedRepLibraries.fasta -rmblast_dir rmblast-2.11.0/bin -crossmatch_dir /sw/bin aPleWal_genome.fasta

### Genome annotation

#### Transcriptome and transcript mapping

cDNA from *de novo* transcriptomes^5, 24^ and high-quality Iso-seq reads from three libraries (brain, spleen and blastema) were aligned to the *Pleurodeles* genome using minimap2 (v 2.24-r1122) and the following arguments:

-I200G -axsplice:hq --secondary=no -uf -G5m -a

Gff files from the resulting sam file were created using the following commands:

samtools view -b in.sam | samtools sort -o in.sorted.bam
bedtools bamtobed -bed12 -i in.sorted.bam > in.bed
bedToGenePred in.bed in.genepred
genePredToGtf \"file\" in.genepred in.gtf
genometools-1.6.1/bin/gt gtf_to_gff3 -tidy in.gtf | genometools-1.6.1/bin/gt gff3 -tidy -sort > out.gff

#### Augustus

The gff files produced from mapping transcripts to the genome (see Transcriptome and transcript mapping section) were used as evidence for augustus (v 3.4.0) with the following arguments:

augustus --extrinsicCfgFile=extrinsic.M.RM.E.W.cfg --hintsfile=hints.gff --species=chiloscyllium pleurodeles.fasta > augustus.predictions.gff

The species *chiloscyllium* was chosen as a model as Braker was not able to produce a *de novo* model for *Pleurodeles* due to the genome size causing errors in the GeneMark step.

#### EvidenceModeler

The gff files produced from the last two sections were given to EvidenceModeler (v 1.1.1) to filter alignments for correct annotations with weights:

OTHER_PREDICITON rna 2

ABINITIO_PREDICTION AUGUSTUS 1

A chunk size of 30Mb and an overlap size of 10Mb were used to avoid the larger introns being removed due to default sizes used by EvidenceModeler.

As EvidenceModeler aims to only produce one isoform per gene, afterwards we included any transcripts that were present in at least two of the transcript evidences (2 transcriptome assemblies and 3 Iso-seq libraries).

#### Gene nomenclature

Combining “high-confidence” transcripts with the output from EvidenceModeler resulted in 164,283 predicted isoforms including 56,783 conserved protein coding isoforms (i.e., with a significant homology hit E-value cutoff 10^-10 with a protein in UniprotKB’sSwiss-prot database). For the purpose of gene nomenclature and downstream analysis, multiple isoforms belonging to the same gene were grouped under the same name using a custom python script (aPleWal.mergeisoforms.py), which took isoforms with consecutive numbers (e.g. gene47, gene48, gene49, gene50 and gene51) and the same homology hit (e.g. XYNB_NEOPA) and gave them the same gene name (e.g. XYNB.1). In the event that multiple genes shared the same UNIPROT ID, each additional gene was given a greater integer suffix (e.g. XYNB.2, etc.). Transcript isoforms are identified with an additional integer suffix after the gene name (e.g. XYNB.1.1, XYNB.1.2, etc.). The outcome of this process created 142,667 gene models of which 35,167 were conserved protein coding genes. Finally ncbi’s table2asn function (https://ftp.ncbi.nlm.nih.gov/asn1-converters/by_program/table2asn/) was used to filter spurious annotations for features such as internal stop-codons or errors in frame resulting in our gff3 file: [aPleWal1.anno.20220803.gff3](https://edmond.mpdl.mpg.de/file.xhtml?fileld=194630&version=DRAFT).

#### Protein-based synteny

Synteny plots between assemblies were created by aligning protein sequences from *Pleurodeles* annotation [aPleWal1.anno.20220803.gff3](https://edmond.mpdl.mpg.de/file.xhtml?fileld=194630&version=DRAFT) against protein sequences downloaded for *Homo sapiens* (grch38), axolotl -*Ambystoma mexicanum*-(AmexT_v47-AmexG_v6.0-DD), *Protopterus annectens* (GCF_019279795.1_PAN1.0), gar - *Lepisosteus oculatus-* (Lepisosteus_oculatus.LepOcu1) and *Xenopus tropicalis* (GCF_000004195.4_UCB_Xtro_10.0) using blastp (v 2.11.0+) and vice-versa only allowing max_target_seqs=1. In instances where the same transcripts were mapped against each other one-to-one, these locations in the two genomes were included in the synteny plots based on locations in the gff files.

#### Annotation v2

A second version gene annotation was generated using purely transcript data:

RNA-seq data from BioProject PRJNA353981 were aligned to the genome using hisat2 (v2..2.1) cDNA and iso-seq were aligned to the genome using minimap2 as above and bam files were filtered using samtools view and the following arguments filtering for high-quality alignment (Q60) and removing any unpaired, secondary, or supplementary alignments:

-F 3844 -q60

The merged bam file was used as input to stringtie (v2.2.1) and coding sequences were identified using TransDecoder (v5.5.0) after first identifying protein homolog regions mapping to the Swissprot and Pfam databases using blastp (v2.11.0) and hmmscan (v3.3.2) respectively.

Annotations with in-frame or early stop codons were removed using ncbi’s table2asn program and then orthologs to known proteins in Swissprot and Pfam were identified by mapping as above and filtering using Trinotate (v3.2.2) resulting in annotation file: [aPleWal.anno.v2.20220926.gff3](https://edmond.mpdl.mpg.de/file.xhtml?fileld=196589&version=DRAFT). This annotation resulted in 65,597 gene models, 174,782 transcripts and 14,752 single exon genes.

#### Transposable elements

Among the 33,742 gene models encoding conserved protein coding genes, 14,943 genes were putative transposable elements by virtue of protein homology with a transposable element component.

#### Current protein-coding gene count

The exclusion of 14,943 putative transposable elements from the 33,742 conserved protein coding genes left 18,799 conserved protein coding genes which is the current protein-coding gene count for *Pleurodeles waltl*.

#### Phylogenetic tree

The one-to-one orthologs identified via best-matches between *Pleurodeles* and axolotl, *Xenopus*, *Gar*, *Danio rerio* (GCF_000002035.6_GRCz11), *Protopterus*, human and *Gallus gallus* (bGalGal1.mat - GCF_016699485.2) as above using blastp on protein sequences from ncbi. Multiple alignments between all orthologs were created using MAFFT (v7.505 https://doi.org/10.1093/nar/gkf436) with arguments –auto. For each ortholog alignment, maximum likelihood trees were created using RAxML (v8.2.12 https://doi.org/10.1093/bioinformatics/btu033) using arguments: -f a -m PROTGAMMAAUTO -p 15256 -T 50 -x 271828 -N 100 -o zebrafish,gar. Finally, a consensus tree was created using MP-EST (v.2.1 https://doi.org/10.1186/1471-2148-10-302) with all 7.524 pairwise ortholog trees.

### Intron counting

Gene intron coordinates were retrieved from the *P. waltl* genome annotation gff-file using faidx from SAMtools^38^ 1.12 and the complement tool from BEDtools^39^ 2.30.0. First, the genome annotation was formatted to gtf with AGAT Toolkit^40^ (agat_convert_sp_gff2gtf.pl). Afterwards, the chromosome sizes were calculated with faidx, and the intergenic regions were retrieved from the annotation file. Finally, the intron coordinates were obtained by crossing the intergenic regions with the exon coordinates from the annotation file with the bedtools complement command.

### Repetitive element annotation and analysis

#### Repeat library assembly

The *de novo* repeat libraries of *A. mexicanum, X. tropicalis, and H. sapiens* genomes were developed via RepeatModeler 2.0.2a^41^, with RECON 1.0.8^42^, RepeatScout 1.0.6^43^, and LTRStruct (LTR_retriever 2.9.0^44^ and LTRharvest^45^ from genometools 1.6.2) methodologies for the identification of repetitive elements. Repetitive elements classified by RepeatModeler as “Unknown” were further processed with the DeepTE^46^ algorithm to identify their possible family.

A *de novo* repeat library of each genome was made into a database with RepeatModeler BuildDatabase function. Then, a fasta file with predicted classified repeats and their sequences was generated:

RepeatModeler-2.0.2a/RepeatModeler -database "REF_db" -pa 32 -LTRStruct

DeepTE was executed with a metazoan repeat training dataset to classify “Unknown” repeats:

python DeepTE/DeepTE.py -d deep_temp -o REF_DeepTE -sp M -m_dir Metazoans_model
-i repeatFamiliesUnknowns.fa

As a final step, the obtained *de novo* library was combined with the Dfam 3.3 database consensus:

RepeatMasker/famdb.py -i RepeatMasker/Libraries/Dfam.h5 families --ancestors -f fasta_name "organism name" > Dfam.fasta
cat Dfam.fasta db-families.fa > repeatFamilies.fasta

#### Masking of transposable elements

To identify the location of the predicted repeats, the combined Dfam and *de novo* library consensus were mapped to the whole genome using RepeatMasker 4.1.2-p1^47^ with RMBlast 2.11.0 as a search engine. This generated a gff-annotation file with repetitive elements position across the genome:

RepeatMasker -pa 32 -gff -xsmall -dir repeat_masker/ -e rmblast
-libdir RepeatMasker/Libraries -lib repeatFamilies.fasta genome.fa

#### Classification procedure of repetitive elements

To assess repeat composition, a custom bash script was written to process files in an automatic manner. Classification of repeats was based on^48^ and was expanded with further information on unclassified ancient repeats from Repbase^49^. The categorization of other non-transposable elements was based on the Dfam classification. Each element was assigned to a class, a superfamily, a family, and a category, as shown in Table 1.

**Table 1.** Classification structure of repetitive elements.

| Class | Superfamily | Family | Category |
| --- | --- | --- | --- |
| Class I | nLTR | R2 | LINE |
| Class I | nLTR | MIR/MIR-core | SINE |
| Class I | LTR | Ty3/Gypsy | LTR |
| Class II | Cryptons | CryptonA/Crypton-A | DNA |
| Class II | DD(E/D) transposons | hAT | DNA |
| Other | Simple_repeats | NA | non-TE |

The generated repeat IDs and their corresponding names from RepeatModeler and DeepTE were combined into a dictionary with added classifications. Files were parsed with GNU Awk, where each string was split to retrieve a family name. This name was then compared with the file with classification structure. In the case of repeats that aligned with the Dfam database consensus, no IDs were generated. Thus, they were processed separately and repeat classification was retrieved from Dfam classification using RepeatMasker inner function famdb.py. The result was a repeat annotation gtf-file with coordinates, the names of repetitive elements, and full classification. This annotation was subsequently used to quantify repeat family contributions to the genome size of *P. waltl*, *A. mexicanum*, *X. tropicalis*, and *H. sapiens*.

#### Genomic location of the annotated repeats

To identify the location of repetitive elements within introns, exons and intergenic regions of the *P. waltl* genome, their coordinates were intersected with bed-files generated in the ‘Intron counting’ section (see above) via BEDtools intersect tool.

### P. waltl transcriptome analysis

For expression analysis, transposons from the hAT family as well as domesticated hATs genes (ZBED) were used. Due to the existence of more than 4 million entries in our hAT annotation file, hAT transposons’ coordinates were split into different files based on their chromosome location and further analysis was done separately. Raw RNASeq data was only used for counting hAT transcripts. The raw counts of ZBED genes were available^5^.

#### Iso-Seq data analysis

Iso-Seq data was mapped to *P. waltl* genome with minimap2 2.24^50^ with following parameters:

minimap2 -I200G -t24 -axsplice:hq --secondary=no -uf -G5m $genome_file $isoseq_file

The resulting sam file was transformed into a bam format, sorted by coordinates and indexed creating a csi-file to accelerate further transcript counting:

samtools view -@24 -b isoseq.pWaltl.mapped.sam | samtools sort -@24 -o isoseq.pWaltl.sorted.bam - && samtools index -c -@24 isoseq.pWaltl.sorted.bam

The number of reads for hAT transposons and ZBED genes was acquired with featureCounts ^51^ from subread 2.0.1 package:

featureCounts -t similarity -g gene_id -L -a annotation.gtf -o $outfile $bam_file

#### RNAseq data analysis

STAR^52^ 2.7.9a package was used to index the *P. waltl* genome and then map RNASeq transcripts to it:

STAR --runThreadN 24 --runMode genomeGenerate --genomeDir ./indexed_genome
--genomeFastaFiles $genome_file --sjdbGTFfile $full_annotation.gtf --sjdbOverhang
124 --limitGenomeGenerateRAM 55000000000

Mapping was performed using parameters previously reported^5^:

STAR --genomeLoad LoadAndRemove --genomeDir $genome_dir --runThreadN 24
--readFilesIn rnaseq1_1.fastq rnaseq1_2.fastq --outFileNamePrefix "rnaseq1_tr_"
--limitGenomeGenerateRAM 300000000000 --runDirPerm All_RWX --outFilterMultimapNmax 1000 -- outFilterMismatchNoverLmax 0.05 --alignIntronMax 1 --alignIntronMin 2
--scoreDelOpen -10000 --scoreInsOpen -10000 --alignEndsType EndToEnd
--limitOutSAMoneReadBytes 100000000000
Sam-file transformations were performed as described for the Iso-Seq analysis. Then transcripts were counted: featureCounts -p -T 10 -t similarity -g gene_id -a $hAT.gtf -o $outfile $bam

#### Data processing

Differential expression of RNASeq data was processed with DESeq2^53^ R package. DESeq2’s median of ratios was used to normalise raw counts to measure up- or downregulation of transposons and genes. Transcriptional count mean-dispersion relationship was performed with parametric fit or local regression fit (transposons). For hATs, analysis was only performed on predicted repeats that were longer than 2000 bp. Samples from 0 days post amputation (dpa) and 3 dpa or 7 dpa were compared pairwise to determine significantly differentially expressed genes/transposons. Genes or repeats were filtered by adjusted p-value (<0.1) for visualisation. After regularised log transformation and data centering Z-scores were calculated and visualised as a heatmap applying pheatmap package. All other plots were generated using package “ggplot2” from “tidyverse”^54^.

### Quantitative RT-PCR

RNA from limb or blastema tissues from 9-month-old newts (6-7cm snout-to-cloaca) was isolated using standard Trizol method as previously described^55^. cDNA was prepared using Superscript-IV kit. qPCR was carried out using iQ SYBR Green supermix (Bio-rad, Hercules, CA) on a Chromo 4 instrument running Opticon 3 software (Bio-rad). Gene expression relative to *actb*2 or *ef1*a was calculated using the Livak 2^−ΔΔCq^ method.

**Table 2.**
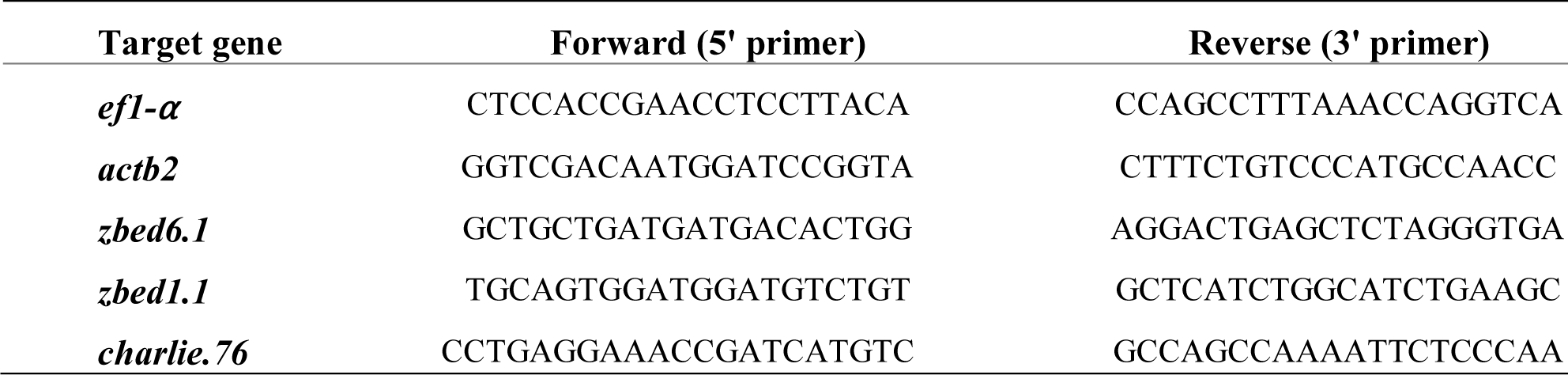
qRT-PCR primers.

## Supporting information

Supplemental material combined

## Acknowledgments

Storage and handling of sequencing data was enabled by resources provided by the Swedish National Infrastructure for Computing (SNIC) at UPPMAX -partially funded by the Swedish Research Council through grant agreement no. 2018-05973-, the DRESDEN Concept Genome Center -part of the technology platform of the CMCB at the TU Dresden, supported by DFG (INST 269/768-1)-, the MPI-CBG computing cloud, and the Center for Information Services and High-Performance Computing (ZIH) at Technische Universität Dresden. Work performed at NGI/Uppsala Genome Center has been funded by RFI/VR and Science for Life Laboratory, Sweden. We thank Miho Kiyooka and Wei Chen for blastema Iso-seq library preparation and Sequel sequencing in the National Institute for Genetics (Japan) and Dominick Kruger, Beate Gruhl and Anja Wagner (CRTD, Germany) for newt husbandry.

## Author contributions

TB performed genome assembly and chromosome scaffolding. AE and TB performed genome annotation, with input and scripts from EO. SI performed computational genomic analysis, with input from MHY, TB, AE, AP and NDL. ES optimized and generated tissue samples for genomic and Iso-seq sequencing. AJA analysed 2n DNA content prior to tissue collection. ES, AJA and NDL coordinated sample extraction. MS, KS, TH, AT generated limb blastema Iso-seq. CO analysed data and performed limb amputations and tissue collection. SI conducted RNA extractions for the limb regeneration curve, primer optimisation and qRT-PCR assays and analysis. TB performed macrosynteny comparisons and haplotype assembly. NDL developed microsynteny pipeline. MHY and AS provided scientific coordination. MHY, AS and NDL supervised the project. AS and MHY provided funding (AS: genome and Iso-seq sequencing except for limb blastema, staff; MHY: computational capacity, staff). AS and MHY edited the manuscript. MHY wrote the manuscript with contributions from all authors.

## Data availability

All data are available in the main text or the supplementary materials. Genome, annotation and haplotype phased assembly files are available through the Max Planck Digital Library at the following location: https://doi.org/10.17617/3.90C1ND and NCBI under the BioProject: PRJNA847026. PacBio HiFi, Hi-C and Iso-seq data will also be available under the same BioProject. Custom annotation scripts are available through https://github.com/egypsci/Pleurodelesgenome.

