## Supplemental material combined for "Sequencing and chromosome-scale assembly of the giant *Pleurodeles waltl* genome"

Brown *et al.*

**This PDF file includes:**

Supplementary Fig. 1 - 15

**Other Supplementary Materials for this manuscript include the following:**

Supplementary data 1 - 7

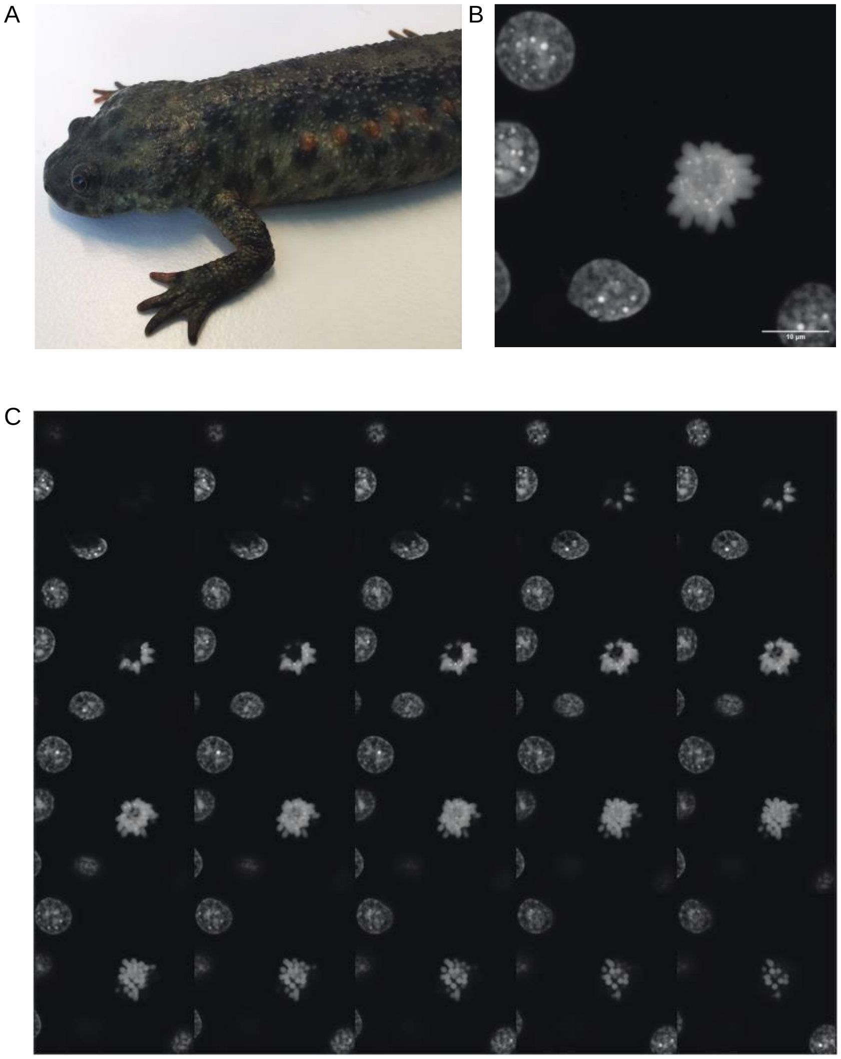


**Supplementary Fig. 1.**

(a) Adult Iberian Ribbed female newt used in the present study for tissue collection and subsequent genome sequencing, Hi-C as well as brain and spleen Iso-Seq. (b) Maximum intensity projection of a Z-stack confocal image of chondrocytes found in the tail vertebra in interphase and metaphase. The cells show both the expected size of the nucleus for a diploid animal (10µm-diameter) and the correct number of chromosomes (2n=24). (c) Individual Z-planes corresponding to Supplementary Fig. 1b.

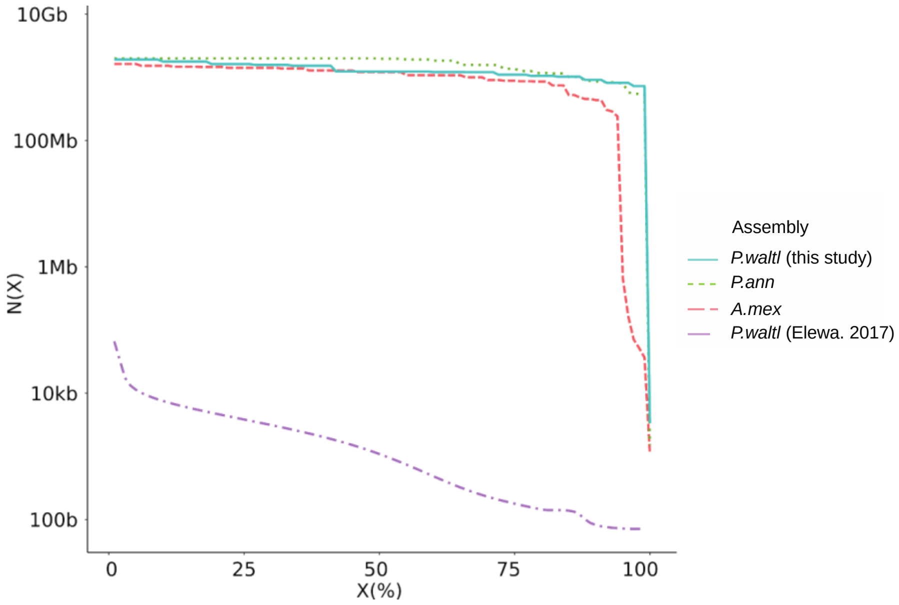


**Supplementary Fig. 2.**

Scaffold N(X) plot showing which % of each assembled genome (X) is contained within pieces at least N(X) bp in size. Shown are contig statistics from *Pleurodeles waltl* (this study and (Elewa et al., 2017)), *Protopterus annectens* (Wang et al., 2021) and *Ambystoma mexicanum* (Schloissnig et al., 2021).

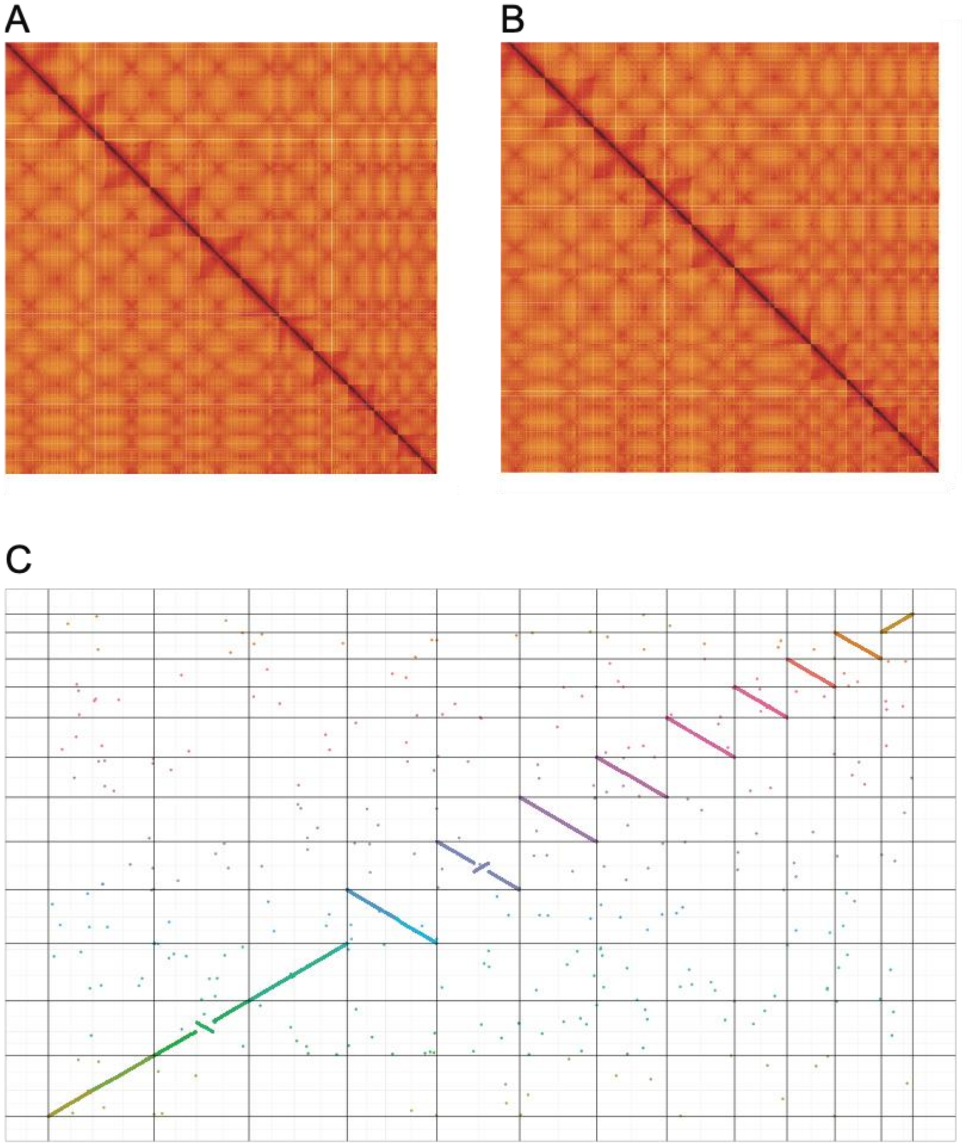


**Supplementary Fig. 3.**

HiC contact maps (a, b) of the two assembled haplotypes and Oxford plot (c) of macrosyntenic relationships between the two assembled haplotypes of *P. waltl.*

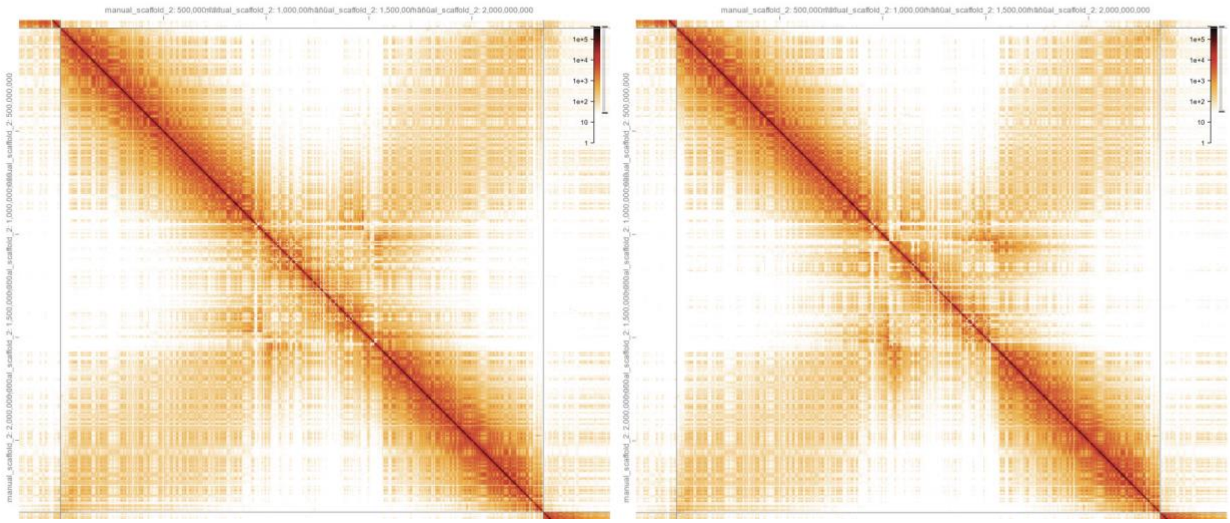


**Supplementary Fig. 4.**

500Mb inversion of the central region of chromosome 2. Hi-C interaction heatmap of contact data for chromosome 2 (left) and corresponding heatmap after inverted central 500Mb region (right).

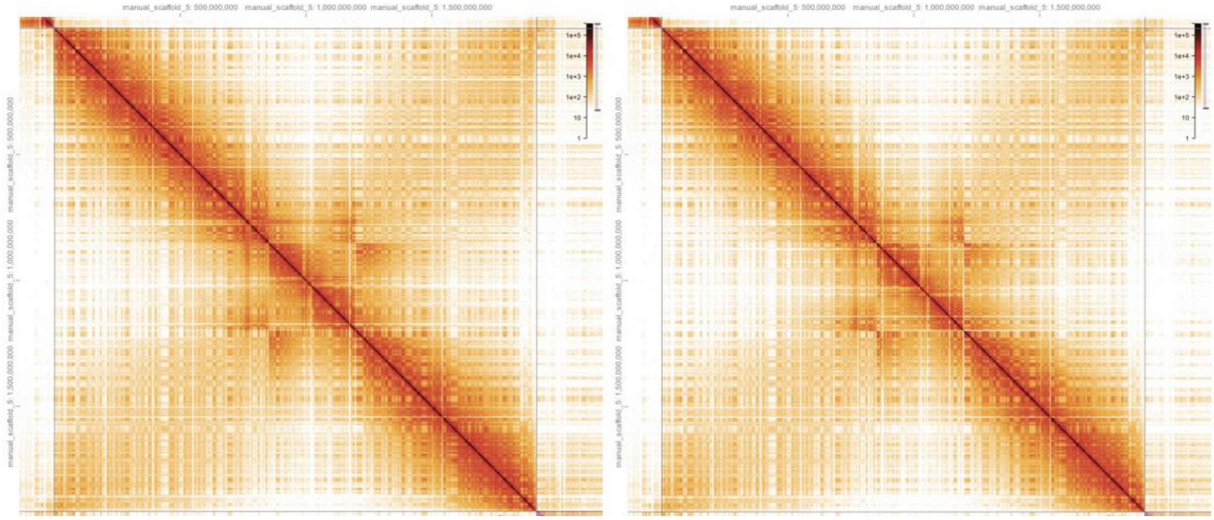


**Supplementary Fig. 5.**

350Mb inversion of the central region of chromosome 5. Hi-C interaction heatmap of contact data for chromosome 5 (left) and corresponding heatmap after inverted central 350Mb region (right).

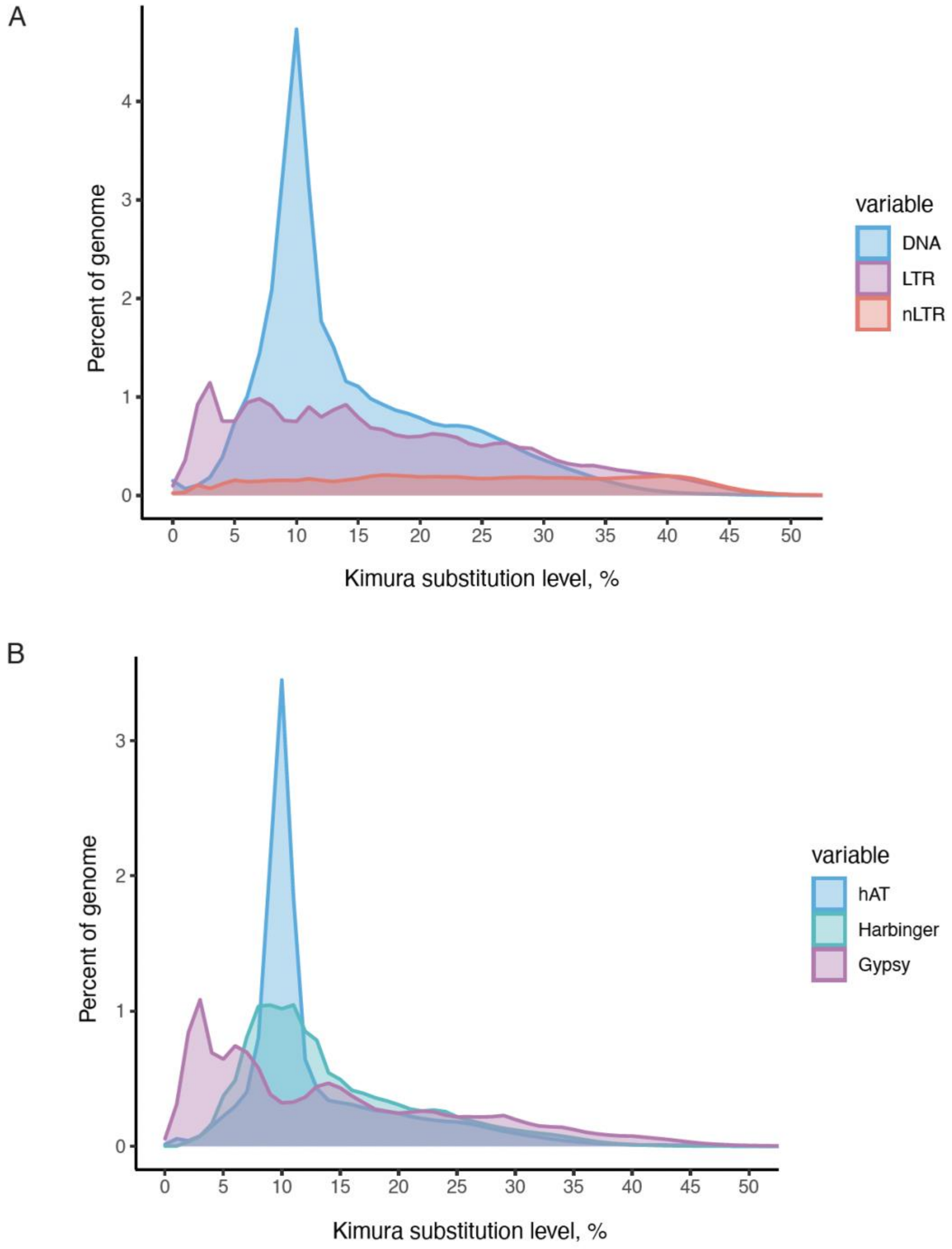


**Supplementary Fig. 6.**

Expansion history of transposable elements based on the kimura substitution level for each copy of the indicated repetitive element against its consensus sequence: (a) for the indicated repeat types; (b) for the top contributor TE families.

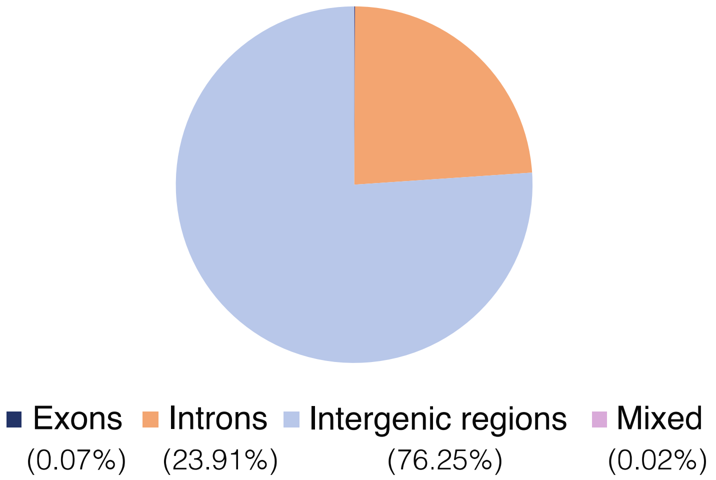


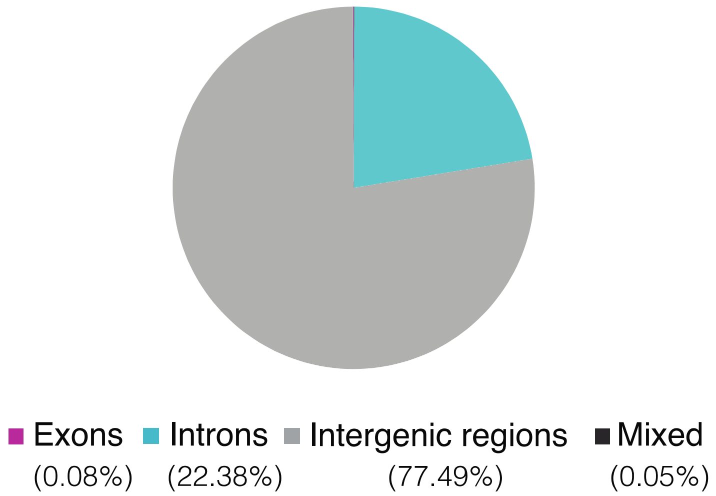


**Supplementary Fig. 7.**

Distribution of hAT elements (top) or all repeats (bottom) within exons, introns and intergenic regions of the *P. waltl* genome, expressed as % contribution to each genomic component.

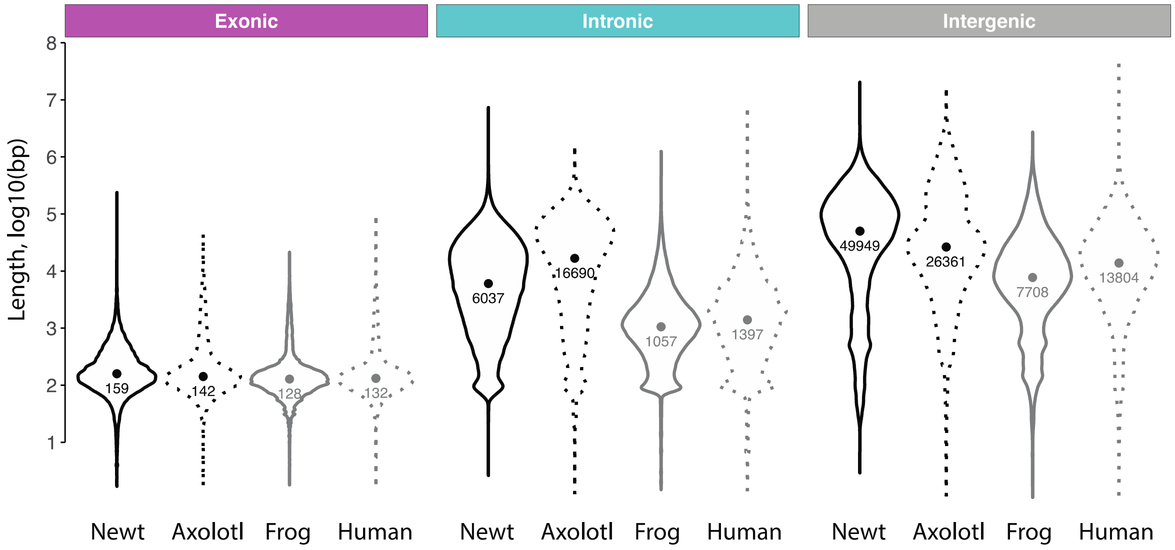


**Supplementary Fig. 8.**

Analysis of exon/intron/intergenic region size in *P. waltl*. Violin plots for intron (left, in kb), intergenic region (middle, in kb) and exon (right, in bp) size. Coloured circles indicate the median of each distribution.

A

blastema

| >2000bp | | | 500-2000bp | | | <500bp | | |
|---|---|---|---|---|---|---|---|---|
| **type** | **number** | **percent** | **type** | **number** | **percent** | **type** | **number** | **percent** |
| Ac | 1 | 0,08 | Ac | 12 | 0,37 | Ac | 78 | 0,91 |
| Charlie | 5 | 0,38 | Charlie | 363 | 11,21 | Charlie | 1027 | 12,01 |
| nMITE | 1273 | 96,66 | nMITE | 2331 | 71,99 | nMITE | 2281 | 26,67 |
| Tip100 | 38 | 2,89 | Tip100 | 4 | 0,12 | Tip100 | 67 | 0,78 |
| | | | MITE | 460 | 14,21 | MITE | 4414 | 51,60 |
| | | | Unknown | 68 | 2,10 | Blackjack | 73 | 0,85 |
| | | | | | | Unknown | 614 | 7,18 |

brain

| >2000bp | | | 500-2000bp | | | <500bp | | |
|---|---|---|---|---|---|---|---|---|
| **type** | **number** | **percent** | **type** | **number** | **percent** | **type** | **number** | **percent** |
| Ac | 1 | 0,27 | | | | Ac | 14 | 0,24 |
| Charlie | 1 | 0,27 | Charlie | 222 | 12,07 | Charlie | 702 | 11,97 |
| nMITE | 355 | 96,99 | nMITE | 1281 | 69,62 | nMITE | 1482 | 25,27 |
| Tip100 | 9 | 2,46 | Tip100 | 2 | 0,11 | Tip100 | 43 | 0,73 |
| | | | MITE | 281 | 15,27 | MITE | 3069 | 52,33 |
| | | | Unknown | 54 | 2,93 | Blackjack | 69 | 1,18 |
| | | | | | | Unknown | 486 | 8,29 |

spleen

| >2000bp | | | 500-2000bp | | | <500bp | | |
|---|---|---|---|---|---|---|---|---|
| **type** | **number** | **percent** | **type** | **number** | **percent** | **type** | **number** | **percent** |
| Ac | 1 | 0,25 | Ac | 2 | 0,10 | Ac | 13 | 0,22 |
| Charlie | 2 | 0,50 | Charlie | 250 | 12,96 | Charlie | 788 | 13,20 |
| nMITE | 383 | 96,23 | nMITE | 1334 | 69,16 | nMITE | 1554 | 26,03 |
| Tip100 | 12 | 3,02 | Tip100 | 4 | 0,21 | Tip100 | 42 | 0,70 |
| | | | MITE | 292 | 15,14 | MITE | 3056 | 51,20 |
| | | | Unknown | 47 | 2,44 | Blackjack | 75 | 1,26 |
| | | | | | | Unknown | 441 | 7,39 |

B

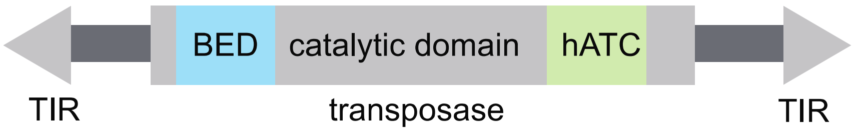


**Supplementary Fig. 9.**

(a) Expression of hAT elements in Iso-seq-derived transcriptomes from limb blastema, brain and spleen according to element length (bp). (b) Schematic diagram representing the structure of a standard hAT, indicating key domains and transposition sites. BED: Zinc Finger BED domain; TIR: Terminal Inverted Repeat; hATC: C terminal dimerization domain.

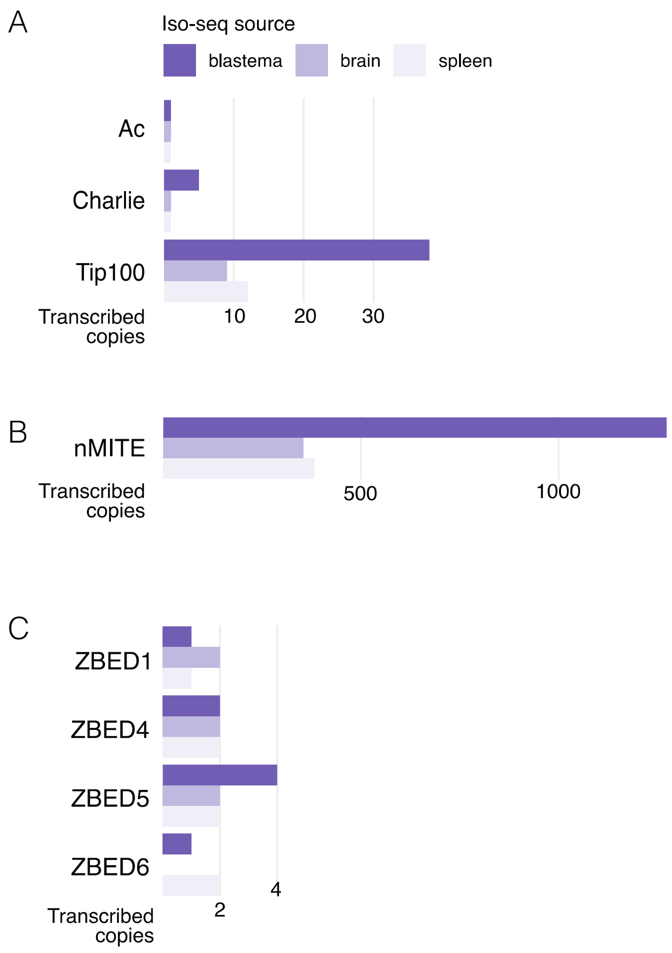


**Supplementary Fig. 10.**

Number of transcribed copies for the indicated hAT (a, b) or domesticated hAT (c) categories based on PacBio Iso-seq data for limb blastema, brain and spleen.

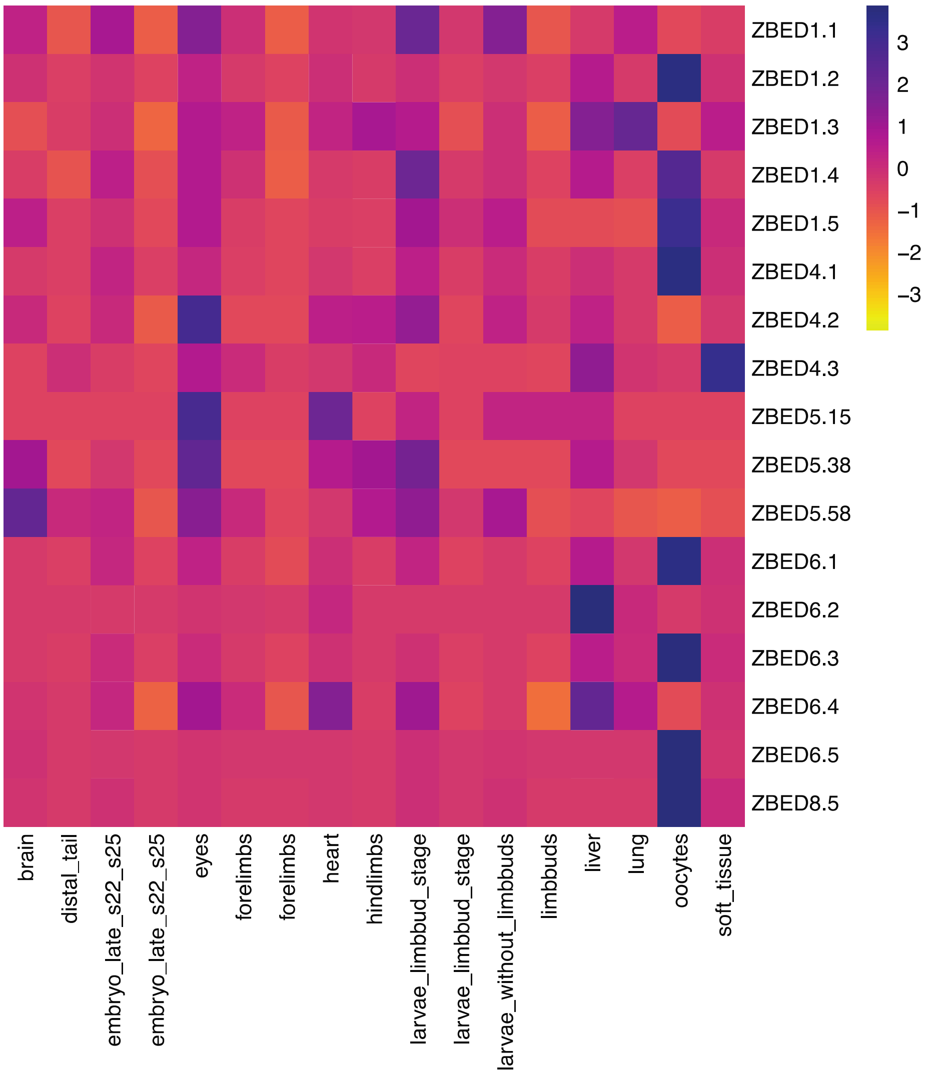


**Supplementary Fig. 11.**

RNAseq quantification of domesticated hATs and relevant isoforms differentially expressed across the indicated *P. waltl* tissues.

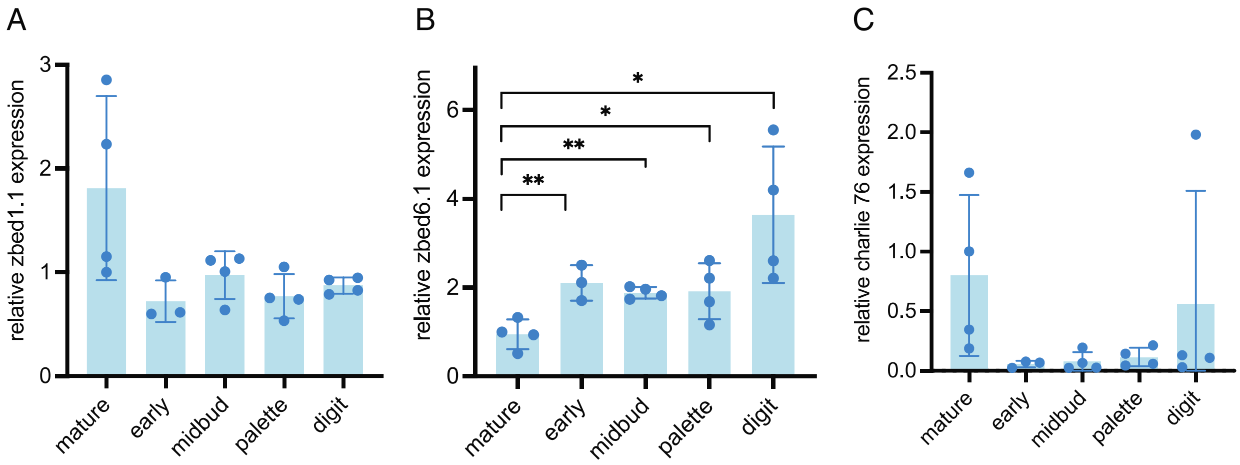


**Supplementary Fig. 12.**

Relative expression of classical and domesticated hATs isoforms during *P. waltl* limb regeneration. qRT-PCR quantification of gene expression for the indicated domesticated (zbed1 (a), zbed6 (b)) and classical hAT (charlie 76 (c)) isoforms relative to *ef1a*. **$p<0.001$ (Welch ANOVA test followed by t-test individual comparison). Error bars represent SD.

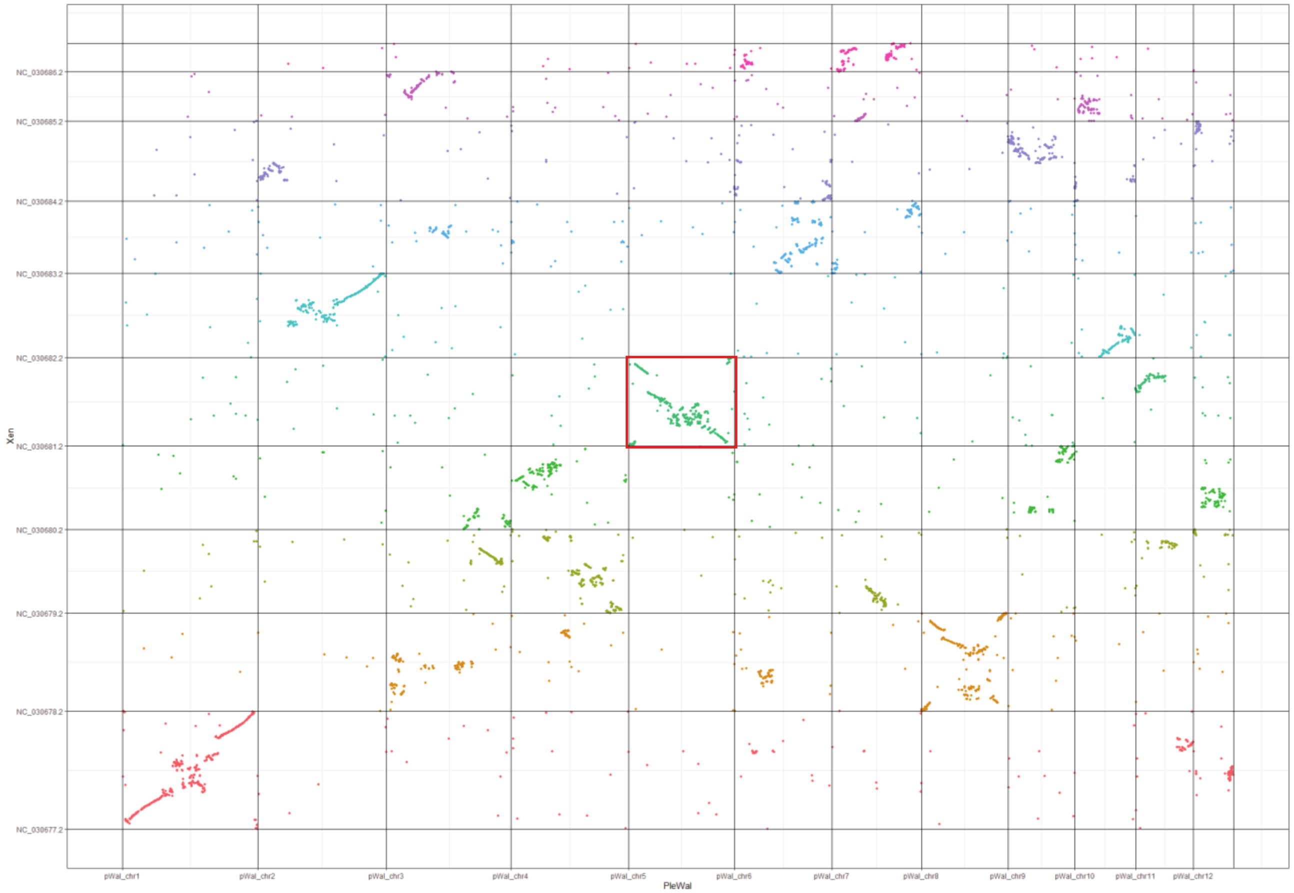


**Supplementary Fig. 13.**

Oxford plot of macrosyntenic relationships between *P. waltl* and *X. tropicalis* chromosomes based on 13,736 one-to-one orthologues. Coloured dots indicate relative chromosomal arrangement of newt-frog orthologues. Inversion of *P. waltl* chromosome 5 is indicated by the red rectangle.

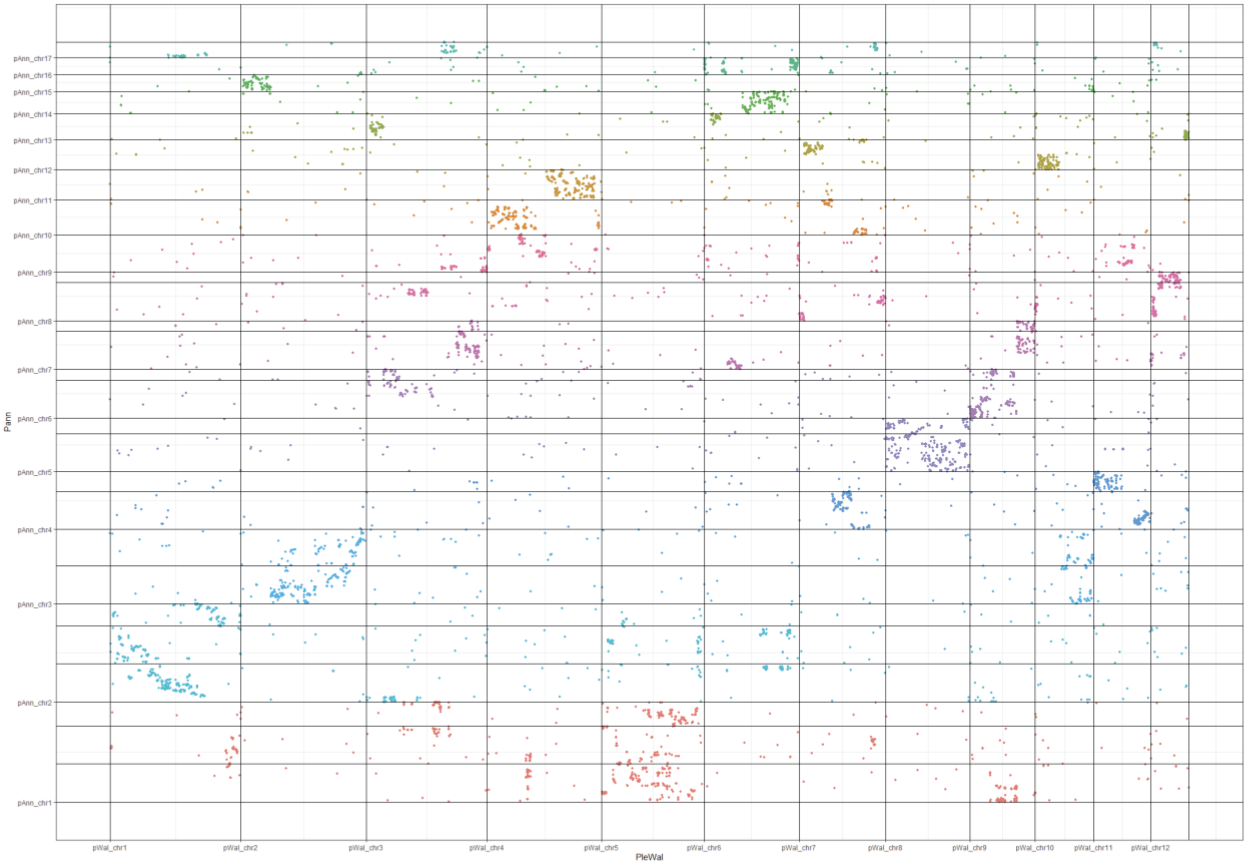


**Supplementary Fig. 14.**

Oxford plot of macrosyntenic relationships between *P. waltl* and *P. annectens* chromosomes based on 13,423 one-to-one orthologues. Coloured dots indicate relative chromosomal arrangement of newt-lungfish orthologues.

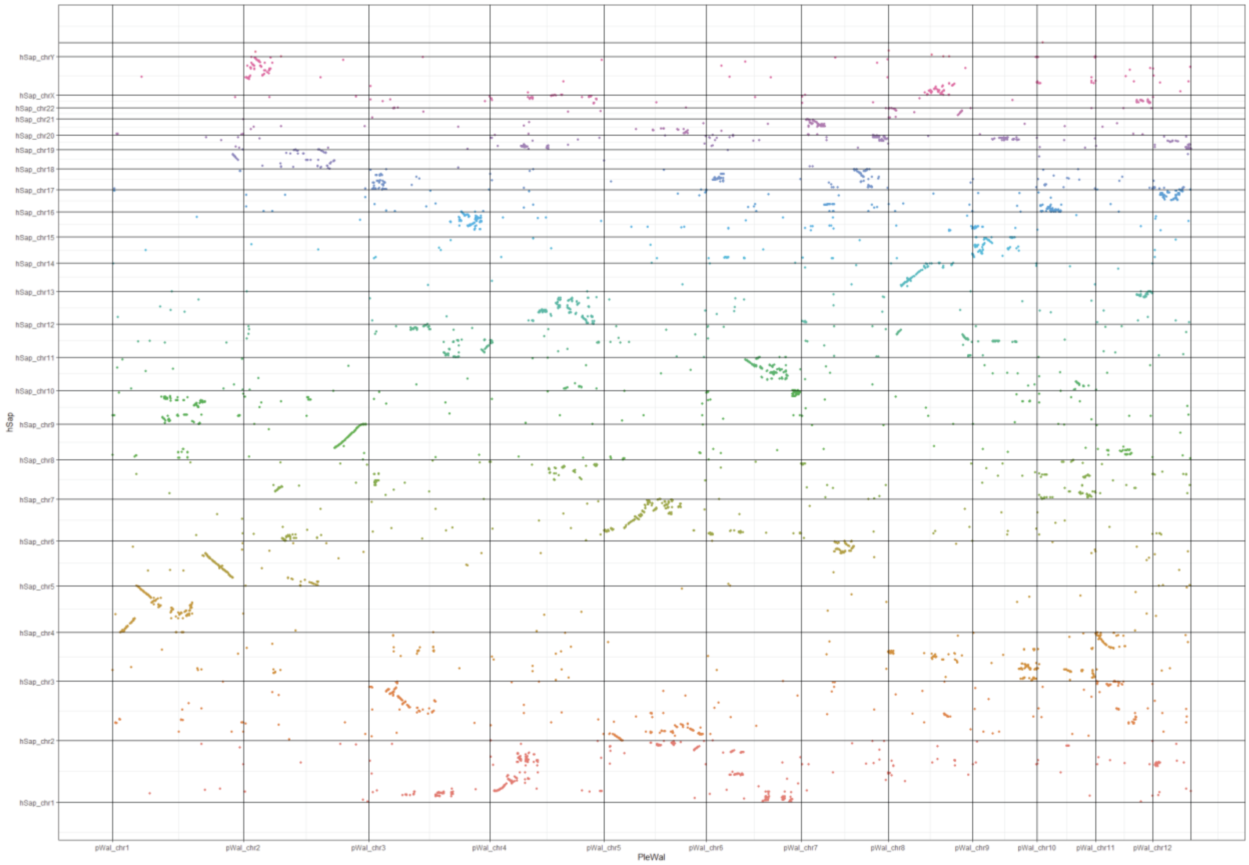


**Supplementary Fig. 15.**

Oxford plot of macrosyntenic relationships between *P. waltl* and *H. sapiens* chromosomes based on 13,127 one-to-one orthologues. Coloured dots indicate relative chromosomal arrangement of newt-human orthologues.

**Supplementary Data (separate files)**

**Supplementary data 1.** GenomeScope profile (k-mer based statistical analysis) of the *P. waltl* genome assembly. (ab) indicates level of heterozygosity; (aa) indicates level of homozygosity.

**Supplementary data 2.** List of genes contained within the 500Mb inverted region of chromosome 2.

**Supplementary data 3.** List of genes contained within the 300Mb inverted region of chromosome 5.

**Supplementary data 4.** DGE analysis of hAT transposons (over 2000bp) during *P. waltl* limb regeneration (0, 3 and 7dpa)

**Supplementary data 5.** DGE analysis of domesticated hAT genes during *P. waltl* limb regeneration (0, 3 and 7dpa)

**Supplementary data 6.** A pipeline for microsynteny analysis.

**Supplementary data 7.** DGE analysis of genes neighbouring the *Tig1* locus during *P. waltl* limb regeneration (0, 3 and 7dpa).
